# Characterization of basal forebrain glutamate neurons suggests a role in control of arousal and avoidance behavior

**DOI:** 10.1101/2020.06.17.157479

**Authors:** James T. McKenna, Chun Yang, Thomas Bellio, Marissa B. Anderson-Chernishof, Mackenzie C. Gamble, Abigail Hulverson, John G. McCoy, Stuart Winston, James M. McNally, Radhika Basheer, Ritchie E. Brown

## Abstract

The basal forebrain (BF) is involved in arousal, attention, and reward processing but the role of individual BF subtypes is still being uncovered. Glutamatergic neurons are the least well-understood of the three major BF neurotransmitter phenotypes. Here we analyzed the distribution, size, calcium-binding protein content and projections of the major group of BF glutamate neurons expressing the vesicular glutamate transporter subtype 2 (vGluT2) and tested the functional effect of activating them. Mice expressing Cre recombinase under control of the vGluT2 promoter were crossed with a reporter strain expressing the red fluorescent protein, tdTomato, to generate vGluT2-cre-tdTomato mice. Immunohistochemical staining for choline acetyltransferase and a cross with mice expressing green fluorescent protein selectively in GABAergic neurons confirmed cholinergic, GABAergic and vGluT2+ neurons represent separate BF subpopulations. Subsets of BF vGluT2+ neurons expressed the calcium binding proteins calbindin or calretinin, suggesting that multiple subtypes of BF vGluT2+ neurons exist. Anterograde tracing using adeno-associated viral vectors expressing channelrhodopsin2- enhanced yellow fluorescent fusion proteins revealed major projections of BF vGluT2+ neurons to neighboring BF cholinergic and parvalbumin neurons, as well as to extra-BF areas involved in the control of arousal and regions responding to aversive or rewarding stimuli such as the lateral habenula and ventral tegmental area. Optogenetic activation of BF vGluT2 neurons in a place preference paradigm elicited a striking avoidance of the area where stimulation was given. Together with previous optogenetic findings suggesting an arousal-promoting role, our findings suggest BF vGluT2 neurons play a dual role in promoting wakefulness and avoidance behavior.

## INTRODUCTION

The basal forebrain (BF) is an important brain region involved in the control of sleep-wake behavior, attention and reward processes (Detari et al., 1999; Zaborszky and Duque, 2003; Brown et al., 2012; Brown and McKenna, 2015; Lin et al., 2015). The BF contains three main populations of neurons utilizing the neurotransmitters, acetylcholine, GABA and glutamate (Gritti et al., 1993, 2006; Hur and Zaborszky, 2005). Of these three neuronal phenotypes, BF glutamatergic neurons are the least well-understood since only recently have tools become available to study these neurons.

The identification of vesicular glutamate transporters (vGluT) as selective markers for glutamate neurons allowed selective identification of glutamatergic somata in BF and throughout the brain using *in situ* hybridization (Herzog et al., 2001; Fremeau et al., 2004; Hur and Zaborszky, 2005; El Mestikawy et al., 2011); as well as BF glutamatergic terminals in projection regions using immunohistochemistry (Henny and Jones, 2006; 2008). vGluT1 and vGluT3 are present at only low levels with BF, with vGluT3 located in a subset of cholinergic neurons which lack the p75 neurotrophin receptor and project to the amygdala (Nickerson Poulin et al., 2006). Thus, vGluT2 is the most prevalent isoform within the BF (Fremeau et al., 2001; Herzog et al., 2001; Hur and Zaborszky, 2005). The development of vGluT2-cre mice (Vong et al., 2011) allowed selective targeting and manipulation of these neurons using double-floxed adeno-associated viral vectors (Anaclet et al., 2015; Xu et al., 2015). Furthermore, crossing vGluT2-cre mice with a cre-dependent reporter strain expressing the red fluorescent protein, tdTomato, facilitated identification of vGluT2 neurons for anatomical and *in vitro* electrophysiological studies (McKenna et al., 2015a, b, 2016; Xu et al., 2015; Yang et al., 2017). Here we used vGluT2-cre-tdTomato mice to precisely characterize the distribution, calcium-binding protein content and projections of BF vGluT2+ neurons. Major subpopulations of BF vGluT2 neurons contained the calcium-binding proteins calbindin or calretinin, suggesting functionally distinct subpopulations.

Analysis of the intra- and extra-BF projections of BF vGluT2+ neurons gives helpful clues to aid us in understanding their functional role. *In vitro* studies in mice suggested excitatory intra-BF connections to PV and somatostatin-containing GABAergic neurons as well as to cholinergic neurons (Xu et al., 2015). Both retrograde (Hur and Zaborszky, 2005) and anterograde studies (Henny and Jones, 2006; 2008; Do et al., 2016; Agostinelli et al., 2017, 2019) in the rodent suggested that subsets of BF vGluT2+ neurons are projection neurons which target the frontal cortex, lateral hypothalamus and other brain areas. Our anterograde tracing experiments demonstrated projections to brain areas involved in the control of arousal and (negative) reward processing. We therefore directly evaluated the role of BF vGluT2 neurons in place preference/avoidance, and report that optogenetic stimulation experiments *in vivo* revealed a dramatic avoidance of the area where stimulation was given. Thus, BF vGluT2 neurons play a dual role in promoting wakefulness and avoidance behavior. Early descriptions of these results were presented previously in abstract form (McKenna et al., 2015a, b, 2016).

## METHODS

### Animals

vGluT2-cre (strain 016963 or congenic strain 028863), PV-cre (congenic strain 017230), ChAT-cre mice (congenic strain 028861) and cre-reporter mice expressing the red fluorescent protein, tdTomato (007905) were purchased from Jackson Labs (Bar Harbor, ME, USA). GAD67-GFP knock-in mice were originally provided by the laboratory of Yuchio Yanagawa (Tamamaki et al., 2003; McKenna et al., 2013) and were generated from our in-house colony. Both male and female animals were used for experiments. Mice were housed under constant temperature and a 12:12 light:dark cycle (7AM:7PM), with food and water available *ad libitum*. All experiments conformed to U.S. Veterans Administration, Harvard University, and U.S. National Institutes of Health guidelines on the ethical use of animals. All measures were taken to minimize the number of animals used and their suffering and were carried out in accordance with the National Institute of Health Guide for the Care and Use of Laboratory Animals (NIH Publications No. 80-23). Experimental procedures were approved by the Institutional Animal Care and Use Committee (IACUC) of the VA Boston Healthcare System.

#### Generation of vGluT2-cre-tdTomato mice

In order to identify BF vGluT2+ (glutamatergic) neurons, we crossed vGluT2-cre mice (mice expressing the enzyme Cre recombinase; Strain 016963, Ai9; Jackson Laboratory, Bar Harbor, ME) with a Cre-reporter strain expressing a red fluorescent marker (tdTomato; Strain 007905; Jackson Laboratory) to generate *vGluT2-cre-tdTomato* mice, in which tdTomato is selectively expressed in vGluT2+ neurons. The selective expression of Cre-driven reporters in vGluT2 neurons in vGluT2-cre mice has been validated in many brain areas (Krenzer et al., 2011; Vong et al., 2011), including BF (Anaclet et al., 2015). Although several Cre-reporter strains are available, we used this particular strain of Jackson Laboratory cre-Tomato mice as reporter, since we used this strain in our previous studies of BF PV neurons using PV-cre mice crossed with these reporter mice (McKenna et al., 2013), allowing direct comparison to the results here.

#### Generation of GAD67-GFP mice

To identify BF GABAergic neurons, we used heterozygous *GAD67-GFP* knock-in mice (Tamamaki et al., 2003; Brown et al., 2008; McKenna et al., 2013). Male, heterozygous GAD67-GFP mice on a Swiss-Webster background were crossed with female wild-type Swiss-Webster mice to generate these animals. In a previous study (McKenna et al., 2013) we confirmed that GFP neurons in BF are GABAergic in these animals, using GABA immunohistochemistry.

#### Generation of vGluT2-tdTomato/GAD67-GFP crossed mice

To test if there was co-localization of tdTomato in GABAergic neurons, we crossed vGluT2-tdTomato animals with GAD67-GFP animals. In these mice, both glutamatergic (red) and GABAergic (green) neurons endogenously fluoresce.

### Target area within the BF

Our target area in the BF for both neuronal characterization and anterograde tracing experiments was the same as in our previous studies (McKenna et al., 2013; Yang et al., 2014). We focused on intermediate areas of BF (substantia innominata (SI), horizontal limb of the diagonal band (HDB), magnocellular preoptic nucleus (MCPO) and ventral pallidum (VP)), where previous studies in the rat (Rye et al., 1984; Gritti et al., 2003; Hur and Zaborszky, 2005; Henny and Jones, 2008) and mouse (Anaclet et al., 2015; Do et al., 2016) found neurons projecting to the neocortex, approximately centered in BF at AP +0.14 mm; ML 1.6 mm; and DV −5.3mm. Our analysis did not include the rostral aspect of BF (medial septum, vertical limb of the diagonal band (MS/DBV)), containing neurons largely projecting to the hippocampus, since previous studies have studied the anatomy and physiology of glutamatergic neurons in this region in detail (Manseau et al., 2005; Huh et al., 2010; Fuhrmann et al., 2015; Leao et al., 2015), or the most caudal aspect of the cholinergic BF which extends into and intermingles with neurons in the globus pallidus and lateral hypothalamus.

### Immunohistochemistry methods

#### 1) Choline acetyltransferase (ChAT)

To determine if tdTomato was located in BF cholinergic neurons and to examine BF vGluT2 input to cholinergic neurons, coronal slices from one well were washed with PBS, placed in a blocking solution (0.5% TX-100 in PBS + 3% normal donkey serum (NDS)), and then incubated in rabbit anti-choline acetyltransferase (ChAT, the synthetic enzyme for acetylcholine) for three days at 4°C (1:150; Cat#AB143, EMD Millipore, Billerica, MA). After incubation, tissue was rinsed and treated for 3 hrs at room temperature (RT, ∼24.4°C) in a secondary antibody coupled to a green fluorophore (1:200; donkey anti-rabbit AF488, Cat#A21206, Thermo Fisher Scientific, Cambridge, MA). For cholinergic neuronal labeling in vGluT2-tdTomato/GAD67-GFP crossed tissue (Fig. 1e) and identification of intra-BF glutamatergic input to cholinergic neurons (Fig. 6a), a blue secondary antibody fluorophore (1:200; donkey anti-rabbit AF350, Cat#A10039, Thermo Fisher Scientific) was used instead.

#### 2) Calbindin (Calb)

For localization of Calb and vGluT2-tdTomato/Calb neurons in BF, slices containing BF from vGluT2-tdTomato mice were washed with PBS, placed in a blocking solution (0.5% TX-100 in PBS + 3% NDS) and then incubated for 2 days at 4**°**C in goat anti-calbindin (1:200; Cat#SC7691; Santa Cruz Biotechnology, Dallas, Texas), followed by 2.5 hours incubation at RT of secondary antibody coupled to a green fluorophore (1:200; donkey anti-goat AF488, Cat#A11055, Thermo Fisher Scientific).

#### 3) Calretinin (Calr)

For localization of Calr and vGluT2-tdTomato/Calr neurons in BF, slices containing BF from vGluT2-tdTomato mice were washed with PBS, placed in a blocking solution (0.5% TX-100 in PBS + 3% NDS), and then incubated for 2 days at 4**°**C in goat anti-calretinin (1:200; Cat#AB1550, EMD Millipore), followed by 2.5 hours incubation (RT) of secondary antibody coupled to a green fluorophore (1:200; donkey anti-goat AF488, Cat#A11055, Thermo Fisher Scientific).

#### 4) Parvalbumin (PV)

For identification of intra-BF glutamatergic input to PV neurons (Fig. 6b), slices containing BF from AAV5-DIO-ChR2-EYFP injected vGluT2-tdTomato mice were washed with PBS, placed in a blocking solution (0.5% TX-100 in PBS + 3% NDS), and then incubated in sheep anti-PV overnight at 4°C (1:150; Cat#AF5058, RnD Systems, Minneapolis, Mn). Tissue was then rinsed and incubated for 3 hrs at RT in appropriate secondary antibody, donkey anti-sheep IgG coupled to a blue secondary antibody fluorophore (1:200; donkey anti-sheep AF350, Cat#A21097, Thermo Fisher Scientific, 4 hr incubation at RT).

#### 5) GFP antibody labeling for amplification

To amplify EYFP signal for anterograde tracing using AAV5-DIO-ChR2-EYFP, tissue was washed with PBS, placed in a blocking solution (0.5% TX-100 in PBS + 3% NDS), incubated in mouse polyclonal anti-GFP primary antibody (1:300; Cat#MAB3580, EMD Millipore) for 3 days at 4**°**C, and secondary antibody (1:500; donkey anti-mouse AF488, Cat#A21202, Thermo Fisher Scientific) overnight at 4°C (fridge). For depiction of intra-BF projections (Fig. 6), tissue was then processed for either ChAT or PV labeling (blue chromophore, described above).

### Validation of primary antibodies used for immunostaining

All primary (Table 1) and secondary (Table 2) antibodies used here have been previously validated and used in peer reviewed publications. Regions expressing ChAT immunoreactivity were similar to those previously reported, including the medial septum/vertical limb of the diagonal band, BF subnuclei as investigated here, striatum, and the pontine tegmental region (Rye et al., 1984; Semba and Fibiger, 1992; Gritti et al., 1993). GFP in GAD67-GFP knock-in mice was expressed throughout many brain regions, similar to that previously reported for GABA/GAD67 immunostaining results, including the cortex, preoptic regions, BF, reticular nucleus of the thalamus, and numerous brainstem regions (Gritti et al., 1993, 2003, 2006; Brown et al., 2008; McKenna et al., 2013). The distribution of Calb and Calr staining was similar to that previously published in the rat and mouse, including the cortex, hippocampus, preoptic regions, BF, and thalamus (Jacobowitz and Winsky, 1991; Rogers and Resibois, 1992; Hof et al., 1999; Zaborszky et al., 1999; Gritti et al., 2003). Secondary antibody-only controls were also performed, omitting the above-listed primary antibodies.

**Table 1.**
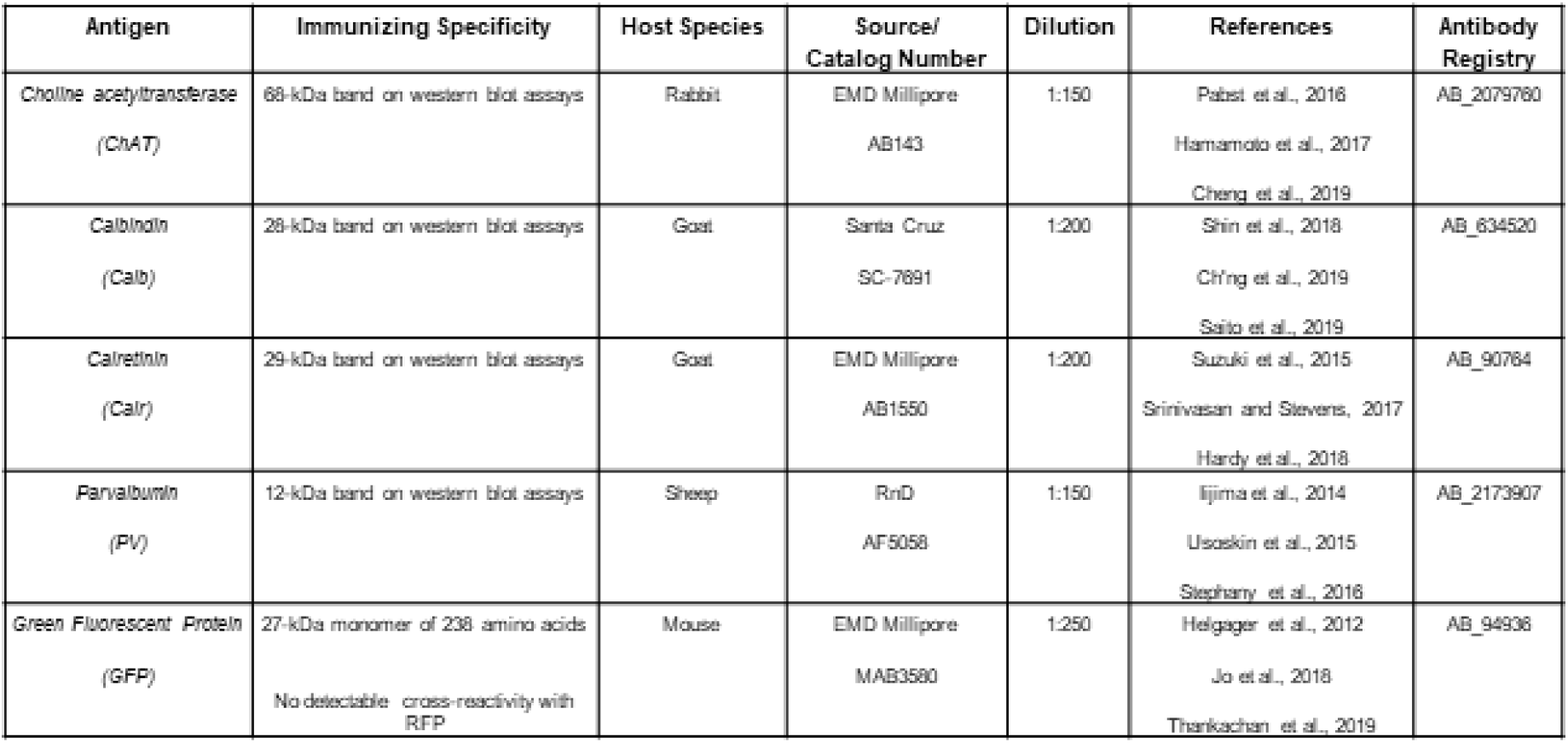
Primary antibody characterization.

| Antigen | Immunizing Specificity | Host Species | Source/<br>Catalog Number | Dilution | References | Antibody<br>Registry |
| --- | --- | --- | --- | --- | --- | --- |
| Choline acetyltransferase<br>(ChAT) | 68-kDa band on western blot assays | Rabbit | EMD Millipore<br>AB143 | 1:150 | Pabst et al., 2016<br>Hamamoto et al., 2017<br>Cheng et al., 2019 | AB_2079760 |
| Calbindin<br>(Calb) | 28-kDa band on western blot assays | Goat | Santa Cruz<br>SC-7691 | 1:200 | Shin et al., 2018<br>Ch'ng et al., 2019<br>Saito et al., 2019 | AB_634520 |
| Calretinin<br>(Calr) | 29-kDa band on western blot assays | Goat | EMD Millipore<br>AB1550 | 1:200 | Suzuki et al., 2015<br>Srinivasan and Stevens, 2017<br>Hardy et al., 2018 | AB_90764 |
| Parvalbumin<br>(PV) | 12-kDa band on western blot assays | Sheep | RnD<br>AF5058 | 1:150 | Iijima et al., 2014<br>Usoskin et al., 2015<br>Stephany et al., 2016 | AB_2173907 |
| Green Fluorescent Protein<br>(GFP) | 27-kDa monomer of 238 amino acids<br>No detectable cross-reactivity with RFP | Mouse | EMD Millipore<br>MAB3580 | 1:250 | Helgager et al., 2012<br>Jo et al., 2018<br>Thankachan et al., 2019 | AB_94936 |

**Table 2.** Secondary antibody characterization.

| **Secondary antibody** | **Vendor** | **Host Species** | **Catalog Number** | **Dilution** | **Incubation** |
| --- | --- | --- | --- | --- | --- |
| *Donkey anti-goat AF488* | Thermo Fisher | Donkey | A11055 | 1:200 | 2.5 hr RT |
| *Donkey anti-mouse AF488* | Thermo Fisher | Donkey | A21202 | 1:200 | 16 hrs 4°C |
| *Donkey anti-rabbit AF350* | Thermo Fisher | Donkey | A10039 | 1:200 | 3 hrs RT |
| *Donkey anti-rabbit AF488* | Thermo Fisher | Donkey | A21206 | 1:200 | 3 hrs RT |
| *Donkey anti-sheep AF350* | Thermo Fisher | Donkey | A21097 | 1:200 | 3 hrs RT |
| *Donkey anti-sheep AF488* | Thermo Fisher | Donkey | A11015 | 1:200 | 3 hrs RT |

### AAV5-DIO-ChR2-EYFP injection for anterograde tracing of BF glutamatergic projections

#### Viral vectors

Viral vectors encoding Channelrhodopsin-2 (ChR2) and enhanced yellow fluorescent fusion proteins (AAV5-DIO-ChR2-EYFP) originally generated by the laboratory of Dr. Karl Deisseroth (Stanford University, CA, USA) were purchased from the University of North Carolina vector core facility.

#### Stereotaxic surgery and viral injection

Mice were deeply anesthetized with isoflurane (1-3%) and viral injections were performed using a 1 µl Hamilton syringe (Cat#7001KH, Point Style 3, Hamilton Company, Reno, NV, USA), targeting BF (AP +0.14 mm; ML 1.6 mm; DV - 5.3mm). 250 nl of viral vector was injected at a flow rate of 25 nl/min. The needle was left in place for an additional 10 min to allow virus diffusion in the brain and to avoid backflow along the needle tract. Once viral injections were complete, the needle was removed. Animals were sacrificed following one month of survival post-injection.

### General immunohistochemistry methods

Mice were deeply anesthetized with sodium pentobarbital (50 mg/ml), exsanguinated with saline and perfused transcardially with a solution of 10 % buffered formalin. Brains were post-fixed for 1 day in 10% formalin, and then transferred to a 30% sucrose solution for 24 hrs at 4°C. Tissue was cut at a thickness of 40 µm on a freezing microtome and collected into four wells of PBS. Following all immunohistochemistry procedures, tissue was washed and mounted onto chrome-alum gelatin-coated slides, dried, and coverslipped using Vectashield hard set mounting medium (H-1400; Vector Laboratories, Inc., Burlingame, CA).

### Chromophore co-localization quantification/photographic depiction

Staining and quantification for neuronal markers/labeling was conducted for each co-localization study. For each immunolabeling stain (ChAT, PV, Calb, Calr) and vGluT2- Tomato/GAD67-GFP crossed animals, representative BF sections [3 coronal section levels per animal (+0.38 mm (Rostral), +.014 mm (Medial), and −0.10 mm (Caudal) from Bregma)] were quantified using Neurolucida software (Version 8; Microbrightfield, Williston, VT) and a Zeiss Imager.M2 microscope outfitted for Structured Illumination (Zeiss Apotome 2). The perimeter, brain areas, and landmarks of BF were first traced at a low magnification (10X), and neuronal location was then plotted at higher magnification (20X). Neuronal markers were then overlayed onto appropriate schematic templates (Franklin and Paxinos, 2008) and replotted using Adobe Illustrator. The distribution of labeled neurons was determined using a mouse brain atlas (Franklin and Paxinos, 2008).

vGluT2-tdTomato cells were identified by the presence of red (excitation/emission at 590:617 nm) fluorescence in the cytoplasm and nucleus. <u>Double-labeled staining of vGluT2-tdTomato tissue:</u> ChAT, Calb or Calr cells were identified by the presence of green fluorescence in the cytoplasm and nucleus (excitation:emission 488:509 nm). <u>vGluT2-tdTomato/GAD67-GFP crossed:</u> GAD67-GFP cells were identified by the presence of green fluorescence in the cytoplasm and nucleus (excitation:emission 488:509 nm). <u>vGluT2-tdTomato mice injected with AAV5-DIO-ChR2-EYFP:</u> EYFP fluorescence was amplified with anti-GFP antibody and appropriate secondary antibody conjugated to a green chromophore (excitation:emission 488:509 nm). ChAT or PV labeled neurons were identified by the presence of blue fluorescence in the cytoplasm and nucleus (excitation:emission 350:450 nm). <u>Long-axis measurement:</u> The long-axis diameter of labeled neurons was measured for each of the 4 BF subnuclei using Neurolucida software. Up to 10 neurons per BF subnuclei (VP, SI, HDB, MCPO) were analyzed, bilaterally, for the 3 representative slices (rostral, intermediate, and caudal) in each animal.

Digital images of fluorescently labeled neurons were captured using a Zeiss Imager.M2 microscope, and select high power photomicrographs were taken using Structured Illumination (Zeiss Apotome 2). Co-localization of select chromophores (e.g., vGluT2-tdTomato and Calb or Calr) was depicted using 3-D Structured Illumination Z-stack photography (Zeiss Apotome 2, Zeiss Imager.M2 system). Structured Illumination photography was also performed to depict BF vGluT2 fibers containing ChR2-EYFP apposed to cholinergic or PV+ neurons, suggesting synaptic contact. Anterograde fiber labeling following AAV5-DIO-ChR2-EYFP injection was evaluated using the Zeiss Imager M2 microscope.

### Statistical Analysis for anatomical experiments

All neuroanatomical density, long-axis diameter, and co-localization measures were analyzed using one-way analysis of variance (ANOVA), in order to compare differences between BF subnuclei (VP, SI, HDB, MCPO), as well as between the three coronal BF slice/section representations (rostral, medial, and caudal). If the main effect of the ANOVA was found to be significant, a pair-wise comparison between groups was then made, using Tukey’s HSD correction. Statistical analysis utilized JMP software (release Pro 14.2.0), and differences determined to be significant when p < 0.05.

### Real-Time Place Preference/Avoidance and Optogenetics

Adult vGluT2-cre, PV-cre or ChAT-cre mice received bilateral injections of 0.7 µl AAV5-DIO-ChR2-EYFP into the BF followed by bilateral implantation of fiber-optic cannulae (200µm, 0.22 NA optical fiber, Doric Lenses at AP 0.0, ML ±1.6, DV −5.4). Each mouse was allowed to recover from surgery for at least 3 weeks prior to experiments.

To test whether optical stimulation affected the preference of the mice for the side of the chamber where optical stimulation was delivered, we used a 3-compartment open chamber (46.5 cm L x 12.7 cm W x 12.5 cm H) consisting of left and right sides (each 17.4 cm L x 12.5 cm H) with different floor patterns and a center compartment (11.7 cm L x 12.5 cm H)(Med Associates, Saint Albans, VT). A camera was placed above the chamber to provide real-time tracking of the position of the mouse in the chamber and to record each trial, for post-hoc analysis using Ethovision XT 14 software (Noldus, Leesburg, VA). Mice were tethered with optical fibers for 5 min before the test. During the test, the mice were released from the center compartment and allowed to freely move between the open chambers for 30 minutes. Whenever the center of the body of the mouse entered one of the two sides, an Arduini Uno with custom code was triggered via a TTL output pulse to continuously drive optical stimulation at frequencies close to the maximal discharge rate of these neuronal types during wakefulness (bilateral, 473 nm, 20 mW, 10 ms pulses, vGluT2: 20Hz, ChAT: 10 Hz, PV: 40Hz) (Xu et al., 2015), with cessation of stimulation when the mouse exited the chamber. The side paired with stimulation (STIM) was counterbalanced between mice. The following day the mice were again placed in the chamber for 15 min without optical stimulation to test if the stimulation resulted in a learned place preference/aversion. Paired t-tests were used to analyze the time spent on the stimulated side vs the unstimulated side.

## RESULTS

Here, we first confirmed that vGluT2-tdTomato neurons within BF represent a separate neuronal population from cholinergic or GABAergic neurons. Their size and relative density within different BF subregions were analyzed together with their coexpression of calcium-binding proteins. Next, we analyzed the intra and extra-BF projections of BF vGluT2 neurons to provide clues to their possible functional role. Analysis of these projections led us to perform optogenetic *in vivo* experiments to test the functional effect of activating them in a place-preference/place aversion paradigm.

### tdTomato is not expressed ectopically in cholinergic or GABAergic neurons in the BF of vGluT2-tdTomato mice (Fig. 1)

To confirm that the fluorescent marker (tdTomato) is not expressed ectopically in BF cholinergic neurons, we performed immunohistochemical staining for the selective marker of cholinergic neurons, ChAT (Fig. 1a, b). ChAT+ and vGluT2-tdTomato neurons were located in all BF subregions. However, none of the ChAT+ cells analyzed in BF (n = 4, 3 sections/animal) co-expressed tdTomato, suggesting that cholinergic neurons do not express vGluT2. To assess co-localization with GABAergic neurons, we crossed vGluT2-tdTomato mice with GAD67-GFP knock-in mice and assessed the extent of co-localization of the red (tdTomato) and green (GFP) fluorescent markers. Again, tdTomato was not expressed in GAD67-GFP (GABAergic) neurons in BF (Fig. 1c), schematically depicted for one representative case in Fig. 1d. These results suggest that neurons with a dual GAD67/vGluT2 phenotype are not present in the BF regions we analyzed. In conclusion, glutamatergic (vGluT2-tdTomato), cholinergic (ChAT), and GABAergic (GAD67-GFP) neurons in BF are three separate neuronal populations in the BF, as illustrated in Figure 1e.

**Figure 1.**
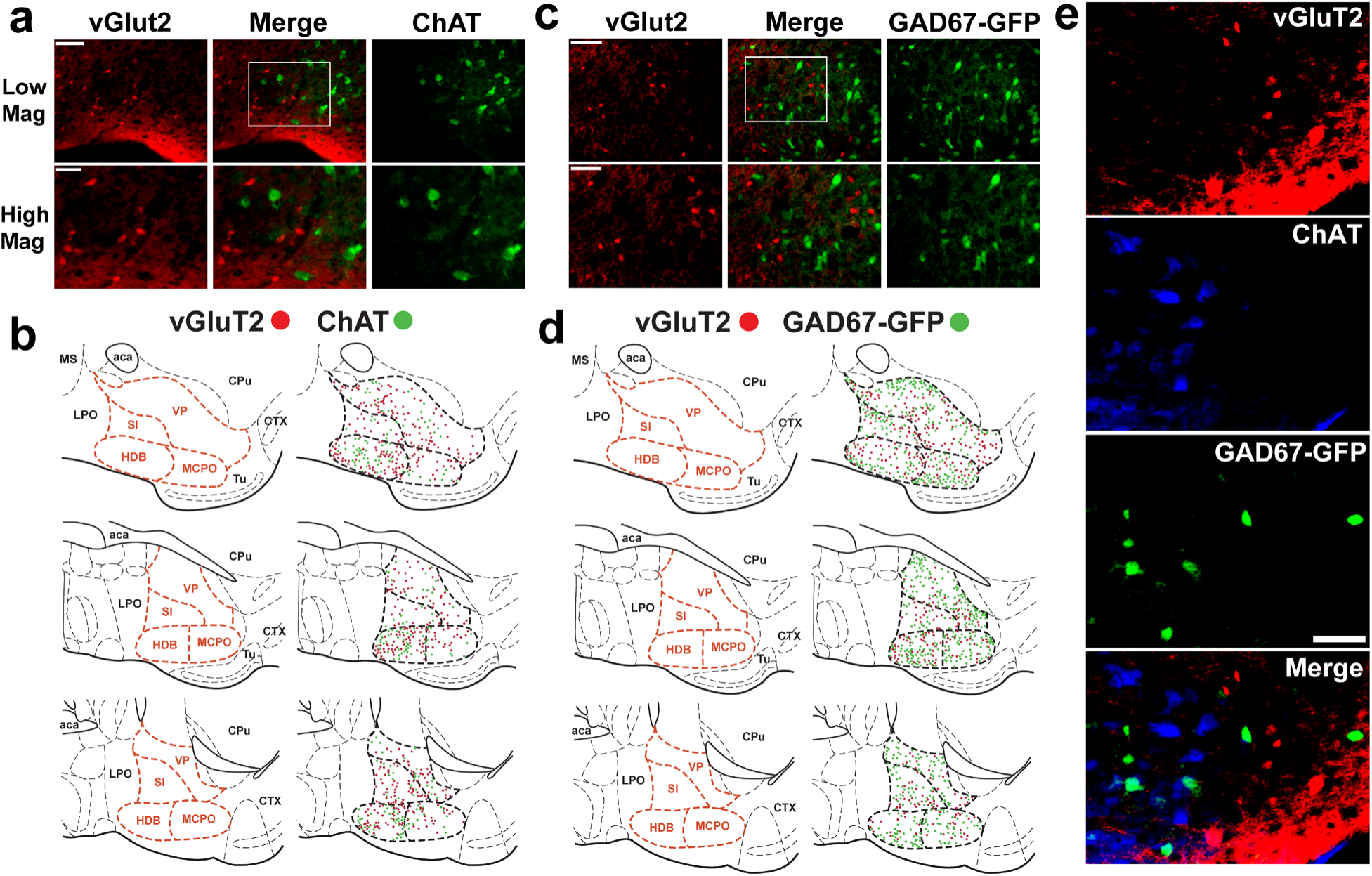
Neurons expressing the vesicular glutamate transporter subtype 2 (vGluT2) are located throughout the basal forebrain (BF) and do not overlap with cholinergic (a-b) or GABAergic (c-d) neurons. a) Fluorescent images of glutamatergic (vGluT2-tdTomato, red) and cholinergic (ChAT, green) neurons in the BF of one representative case, demonstrating lack of co-localization of vGluT2-tdTomato (Red) and ChAT (green) neurons. b) Mapping of the non-overlapping location of glutamatergic (vGluT2-tdTomato, red) and cholinergic (ChAT, green) neurons in intermediate regions of BF. c) Fluorescent images of glutamatergic (vGluT2- tdTomato, red) and GABAergic (GAD67-GFP, green) neurons in the BF of one representative case, demonstrating lack of co-localization. d) Mapping of the non-overlapping location of glutamatergic (vGluT2-tdTomato, red) and more numerous GABAergic (GAD67-GFP, green) neurons in BF. e) Tri-color photographic depiction of vGluT2-tdTomato (glutamatergic, red), ChAT (cholinergic, blue) and GAD67-GFP (GABAergic, green) neurons in BF demonstrates three largely separate neuronal populations in the BF. Scale bars = 100 μm for a and c, 50 μm for e.

### Distribution and long-axis diameter of vGluT2-tdTomato neurons in BF (Fig. 2)

A precise mapping of the distribution, relative density and size of vGluT2 neurons within the mouse BF is not available. Thus, we first quantified the relative density and size (long-axis diameter) of vGluT2-tdTomato neurons in 3 coronal sections spanning intermediate regions of the BF (rostral, medial and caudal slice/section representations), and across the individual subnuclei (VP, SI, HDB, MCPO) using the same methodology that we previously used to map BF GABAergic and parvalbumin neurons (McKenna et al., 2013). vGluT2-tdTomato neurons were localized throughout BF (Figs. 1, 2; n = 4), with the highest density of vGluT2-tdTomato neurons located in MCPO (56.8 ± 3.5 tdTomato+ neurons/mm2/slice; Fig. 2a), followed by the HDB (47.0 ± 3.2) and SI (45.4 ± 5.0) regions. The VP region in the dorsal BF had the lowest densities (25.2 ± 6.4). Significant differences were noted between the four BF subnuclei (F_3,36_ = 5.031, P = 0.005), and post-hoc pairwise comparisons indicated significant density differences between VP vs. HDB (t = 3.00, P = 0.022) and vs. MCPO (t = 3.54, P = 0.005). vGluT2-tdTomato neurons were most dense in the medial portion of BF (46.3 ± 5.0), followed by caudal (36.6 ± 8.0) and rostral (31.9 ± 0.6) regions. To summarize, vGluT2-tdTomato neurons were distributed throughout BF subnuclei, with a decrease in density noted in dorsal BF (VP) when compared to the other subnuclei. Thus, we focused on ventral BF regions in medial BF for our subsequent injections of adenoviral vectors expressing channelrhodopsin2-enhanced yellow fluorescent protein (AAV5-DIO-ChR2-EYFP) for anterograde tracing and *in vivo* optogenetic stimulation experiments.

**Figure 2.**
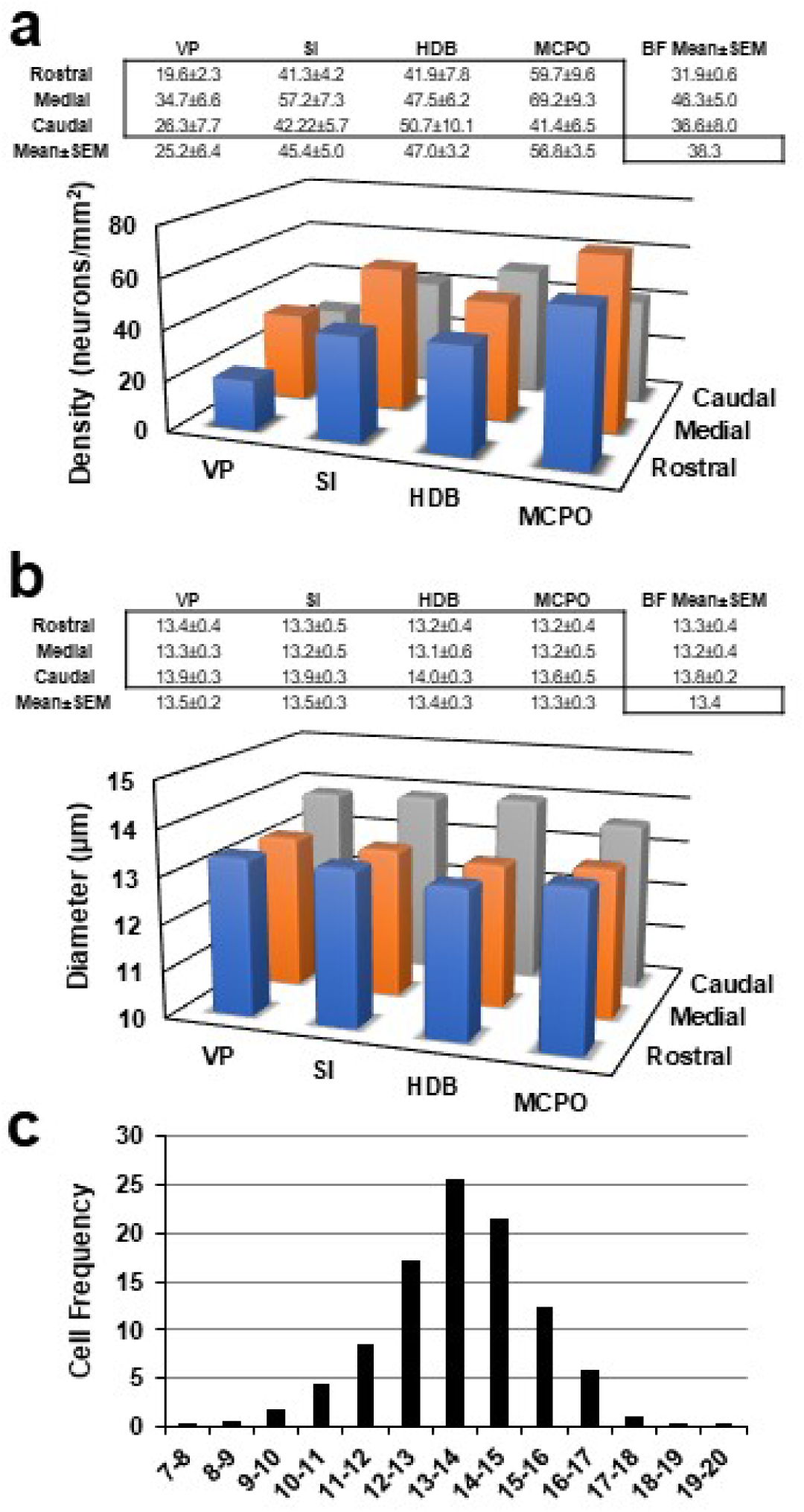
Quantification of the density and size of vGluT2-tdTomato (glutamatergic) neurons in BF subnuclei of vGluT2-tdTomato mice. Measures of the density (a) and long-axis diameter (b) of vGluT2-tdTomato neurons were determined in the ventral pallidum (VP), substantia innominata (SI), horizontal limb of the diagonal band (HDB), and magnocellular preoptic (MCPO) regions of rostral, medial, and caudal BF slices. a) The relative density of vGluT2-tdTomato neurons was highest in the magnocellular preoptic area (MCPO), followed by HDB and SI in the BF of vGluT2-tdTomato mice. Significantly lower densities were found in the VP compared to other BF subnuclei. b) No significant differences were found between mean long-axis diameters of different subnuclei. Comparisons along the rostrocaudal axis revealed that diameters were slightly increased in caudal slices. c) Histogram illustrating the cell frequency distribution for the long-axis diameters of vGluT2-tdTomato neurons (out of 1543 neurons evaluated across four animals), centered around a mean of 13.4 μm. Rostral representation = +0.38 mm from bregma; Medial representation = +0.14 mm from bregma; Caudal representation = −0.10 mm from bregma.

In marked contrast to BF cholinergic, GABAergic and parvalbumin neurons (McKenna et al., 2013), there were virtually no large diameter (>20 µm) BF vGluT2-tdTomato neurons (Fig. 2b). BF vGluT2-tdTomato neurons were almost all small (<15 µm) or medium sized (15-20 µm) neurons (Fig. 2b and c). No significant differences were noted of the vGluT2-tdTomato neuronal long-axis diameters between BF subnuclei (n = 6; VP, 13.5 ± 0.2 µm, SI, 13.5 ± 0.3; HDB, 13.4 ± 0.3; MCPO, 13.3 ± 0.3). Slight differences were noted across the rostrocaudal axis (F_2,36_ = 4.177, P = 0.02; Rostral, 13.3 ± 0.4 µm; Medial, 13.2 ± 0.4 µm; Caudal, 13.8 ± 0.2 µm). Post-hoc analysis revealed that mean values were significantly higher in the Caudal vs. Medial slice (t = 2.63, P = 0.029).

### The calcium-binding proteins calbindin (Calb) and calretinin (Calr) are expressed in subsets of BF vGluT2-tdTomato neurons

Previous studies in the rat demonstrated the presence of calbindin (Calb) in a subset of BF non-cholinergic, non-GABAergic neurons that project to the cortex (Gritti et al., 2003). To determine if Calb labels a subpopulation of BF vGluT2 neurons in the mouse, we performed immunohistochemistry for Calb in vGluT2-tdTomato mice tissue. Indeed, a subpopulation of vGluT2-tdTomato neurons in the BF was Calb (Fig. 3). Photographic depiction (Fig. 3a) and neuronal mapping of one representative case across the rostral-caudal extent of BF (Fig. 3b) demonstrated vGluT2-tdTomato co-localization in a subpopulation of Calb neurons in all subnuclei of BF. Overall, 19.3% of vGluT2-tdTomato neurons also contained Calb in BF (Fig. 3c). Highest levels of co-localization were seen in MCPO (24.8 ± 3.2%), followed by HDB (19.8 ± 3.0%), VP (16.3 ± 1.6%) and SI (16.1 ± 1.2%). Statistical analysis revealed significant differences of co-localization between the BF subnuclei (F_3,36_ = 5.366, P = 0.003). Post-hoc analysis confirmed significant pairwise differences between MCPO vs. VP (t = 3.41, P = 0.008) and vs. SI (t = 3.51, P = 0.006). Significant differences were not found along the rostrocaudal axis.

**Figure 3.**
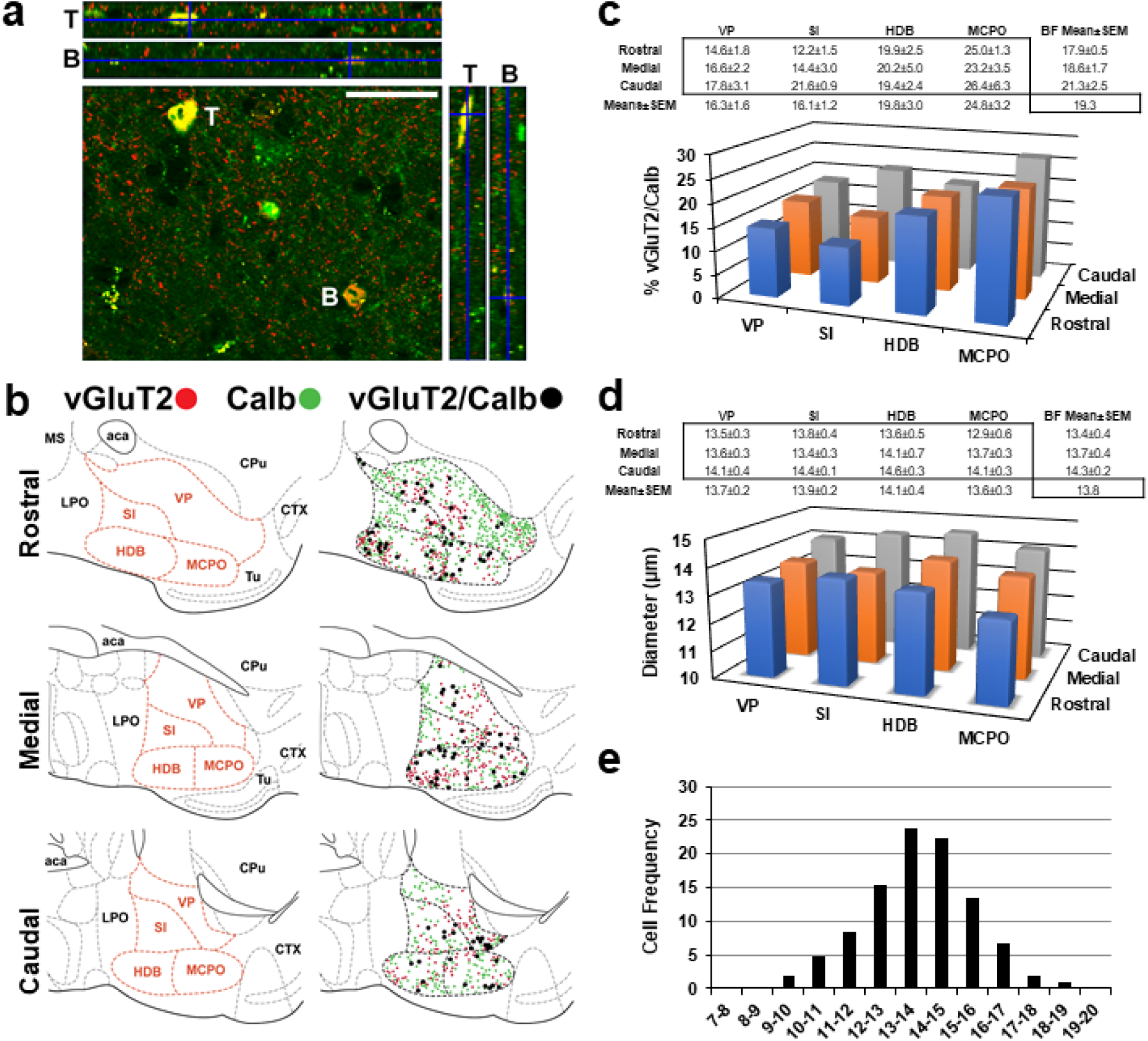
A subpopulation of BF vGluT2-tdTomato neurons co-express the calcium-binding protein calbindin (Calb). a) Fluorescent images (Apotome Structured Illumination, z-Stack, 2 μm thick slices) of glutamatergic (vGluT2-tdTomato, red) and Calb (green) neurons in the BF, demonstrating co-localization. Scale bar = 25 μm. T, top example; B, bottom example. b) Mapping of glutamatergic (vGluT2-tdTomato, red) and Calb (green) neurons depicts many neurons which co-express both markers (black) in BF. c) The highest colocalization was observed in MCPO. There were no significant differences across the rostrocaudal axis. d) The diameter of vGluT2-tdTomato neurons that co-labeled with Calb did not significantly differ between BF subnuclei but caudal vGluT2-tdTomato/Calb neurons were larger compared to rostral and medial slices. e) Histogram illustrating the cell frequency distribution for the long-axis diameters of vGluT2-tdTomato/Calb neurons (out of 834 neurons evaluated across four animals), centered around a mean of 13.8 μm. Rostral representation = +0.38 mm from bregma; Medial representation = +0.14 mm from bregma; Caudal representation = −0.10 mm from bregma.

Measures of the long-axis diameter of vGluT2-tdTomato neurons co-localized with Calb (Fig. 3d) did not differ between BF subnuclei (n = 4; VP 13.7 ± 0.2 µm, SI 13.9 ± 0.2; HDB 14.1 ± 0.4; MCPO 13.6 ± 0.3), but diameter measures did significantly differ between slice representations (F_2,36_ = 4.926, P = 0.012; Rostral 13.4 ± 0.4 µm; Medial 13.7 ± 0.4 µm; Caudal 14.3 ± 0.2 µm), and post-hoc analysis revealed that mean values were significantly higher in the Caudal vs. Rostral (t = 3.07, P = 0.01).The vast majority of vGluT2-tdTomato/Calb neurons were small in size, indicated in the histogram profile (Fig. 3e), centered around the overall size mean of 13.8 µm.

We next performed immunohistochemistry for another calcium binding protein, calretinin (Calr) in the BF of vGluT2-tdTomato mice (Fig. 4). Calr was co-expressed in a subpopulation of vGluT2-tdTomato neurons in all subnuclei of BF (Fig. 4a-b). Overall, 24.6% of vGluT2-tdTomato neurons co-localized with Calr in BF (Fig. 4c; n = 4). Highest levels of co-localization were seen in VP (28.2 ± 1.0%), followed by MCPO (23.6 ± 2.4%), SI (22.5 ± 2.3%) and HDB (21.8 ± 1.3%). Statistical analysis revealed significant differences of co-localization between the BF subnuclei (F_3,36_ = 2.852, P = 0.049). Post-hoc analysis confirmed significant pairwise differences between VP vs. HDB (t = 2.84, P = 0.034). Significant differences were not noted along the rostrocaudal axis (Rostral 24.0 ± 2.8%; Medial 25.2 ± 0.7; Caudal 24.5 ± 2.7). Measures of the long-axis diameter did not differ between BF subnuclei (Fig. 4d; n = 4; VP, 13.9 ± 0.4 µm; SI, 13.6 ± 0.5; HDB, 13.9 ± 0.5; MCPO, 13.5 ± 0.5) or between slice representations (Rostral 13.8 ± 0.5 µm; Medial, 13.1 ± 0.8; Caudal, 14.0 ± 0.4). The vast majority of vGluT2- tdTomato/Calr neurons in BF were small, as indicated by the histogram profile (Fig. 4e), centered around the overall mean of 13.6 µm.

**Figure 4.**
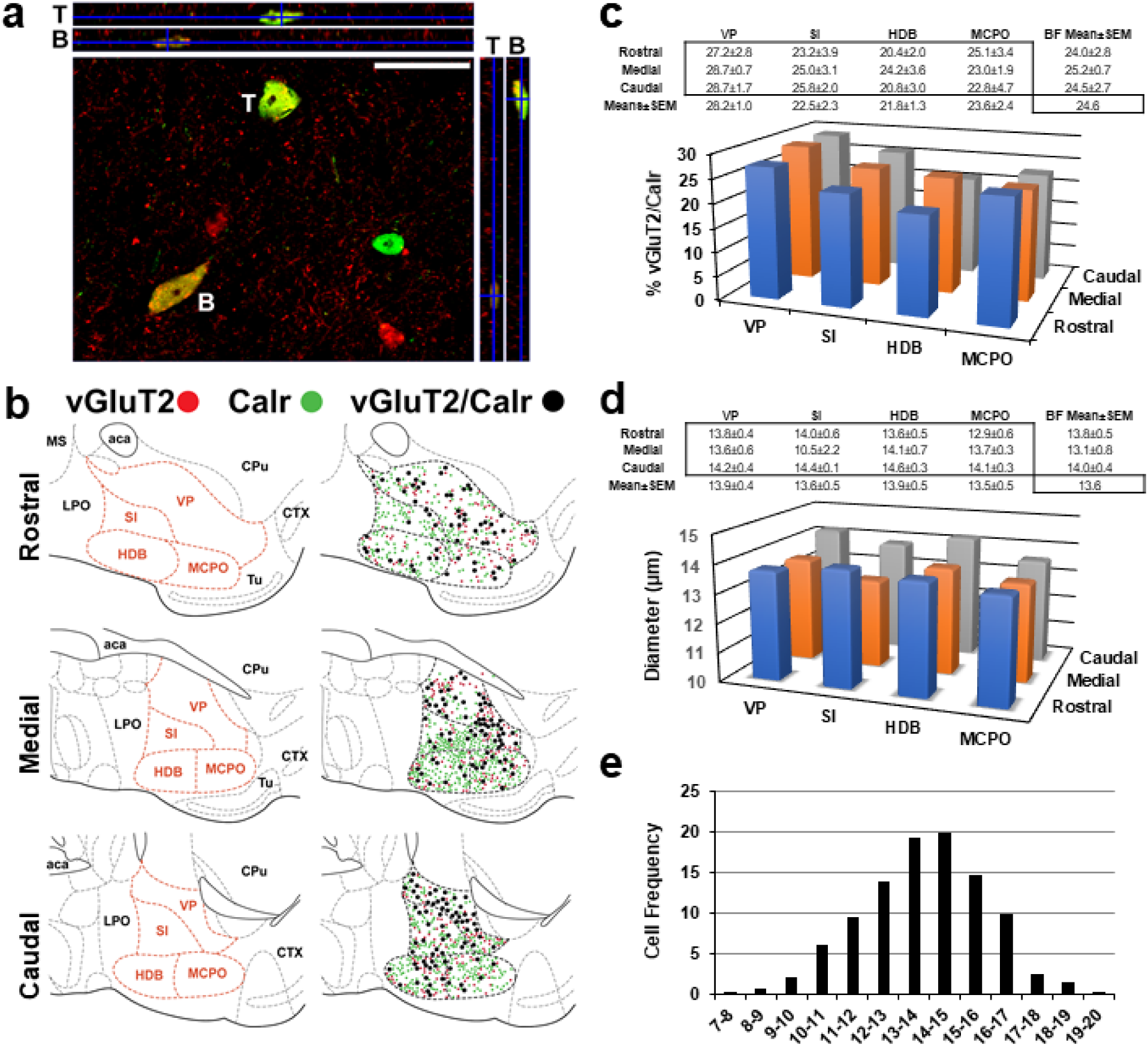
A subpopulation of BF vGluT2 neurons co-express the calcium-binding protein calretinin (Calr). a) Fluorescent images (Apotome Structured Illumination, z-Stack, 2 μm thick slices) of glutamatergic (vGluT2-tdTomato, red) and Calr (green) neurons in the BF, demonstrating co-localization. Scale bar = 25 μm. T, top example; B, bottom example. b) Mapping of glutamatergic (vGluT2-tdTomato, red) and Calr (green) neurons depicts many neurons which co-express both markers (black) in BF. c) Significant differences between BF subnuclei were demonstrated, and the highest colocalization was observed in VP. Significant differences between rostrocaudal slices were not seen. d) The diameter of vGluT2-tdTomato neurons that co-labeled with Calr did not significantly differ between BF subnuclei. e) Histogram illustrating the cell frequency distribution for the long-axis diameters of vGluT2-tdTomato/Calr neurons (out of 1029 neurons evaluated across four animals), centered around a mean of 13.6 μm. Rostral representation = +0.38 mm from bregma; Medial representation = +0.14 mm from bregma; Caudal representation = −0.10 mm from bregma.

Finally, we performed staining for the calcium-binding protein parvalbumin (PV). Only 1.69 ± 0.48 % of BF vGluT2-tdTomato neurons colocalized with parvalbumin; and of all BF PV neurons, 1.46 ± 0.44 % were vGluT2-tdTomato. Thus, we did not perform more detailed analysis of the subregional density or size of these neurons. We previously found that most BF PV neurons are GABAergic (McKenna et al., 2013).

### Anterograde tracing using AAV5-DIO-ChR2-EYFP injections reveals projections of BF vGluT2-tdTomato neurons to neurons/brain areas involved in the control of arousal and the response to aversive and rewarding stimuli

In order to understand the intra- and extra-BF projections of BF vGluT2+ neurons, we performed anterograde tracing experiments using unilateral injections of AAV5-DIO-ChR2-EYFP into the BF of 6 adult vGluT2-cre-tdTomato mice to specifically express ChR2-EYFP in BF vGluT2 neurons. Following a survival period of one month, mice were sacrificed, and brain tissue prepared for subsequent analysis. ChR2-EYFP fusion proteins were expressed in the somata of many vGluT2-tdTomato neurons within the BF (Fig. 5a). >80% of cell somata transduced with AAV co-localized with vGluT2-tdTomato signal, confirming selectivity of viral transduction. Transduced neurons and fibers were located in the target region of BF, indicating successful AAV injection targeting (Fig. 5b). In 4/6 cases transduced neurons were restricted to the BF whereas in 2/6 cases the injections impinged on the neighboring lateral preoptic area (LPO) region (Fig. 5c).

**Figure 5.**
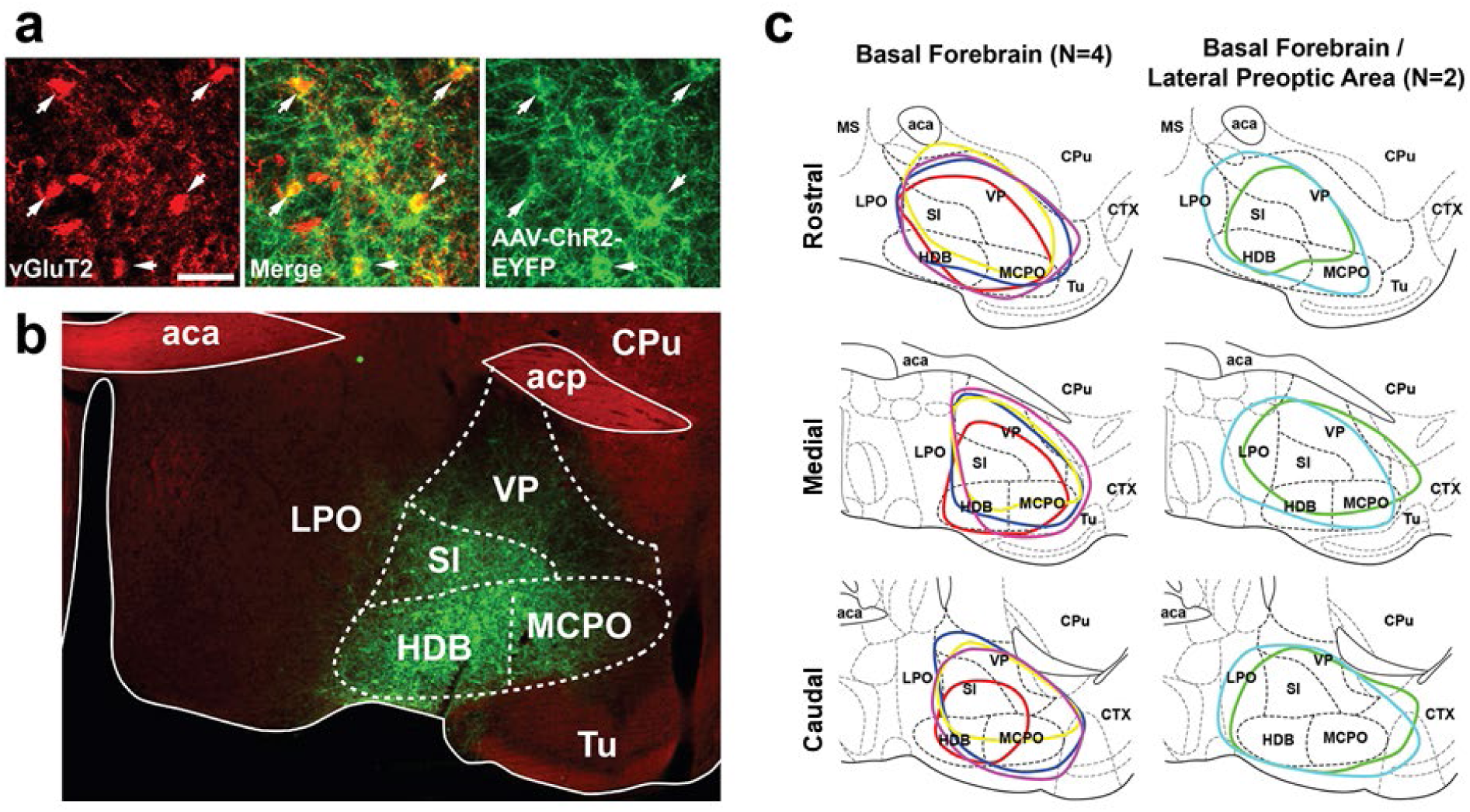
Viral transduction of BF vGluT2 neurons in vGluT2-cre-tdTomato mice. Successful AAV5-DIO-ChR2-EYFP transduction revealed by fluorescence microscopy in vGluT2-cre-tdTomato mice. a) Virally-transduced neurons (EYFP, green) co-localized (arrow) with vGluT2-tdTomato signal (red) in vGluT2-cre-tdTomato mice. Scale bar = 25μm. b) The transduced (AAV5-DIO-ChR2-EYFP, green) neurons were located in the target BF region between −0.10 to 0.38 mm from Bregma (Franklin and Paxinos, 2008). c) Distribution of BF injection sites, represented across three rostral-caudal slices of BF. 2 of the 6 cases impinged on the neighboring lateral preoptic area (LPO). Lower panel shows the location of the cannula tip used for viral injection (coronal sections). Abbreviations: aca, anterior commissure (anterior); acp, anterior commissure (posterior); Cpu, caudate putamen; CTX, cortex; HDB, horizontal limb of the diagonal band; LPO, lateral preoptic area; MCPO, magnocellular preoptic nucleus; MS, medial septum; SI, substantia innominata; Tu, olfactory tubercle; VP, ventral pallidum. Scale bar for a) = 25 µm; b) and c) = 1mm.

We first analyzed the fibers of vGluT2 neurons within the transduced area of BF. As shown in Fig. 6, fibers amplified with anti-GFP antibody (green) closely apposed neurons labeled with ChAT (indicating cholinergic phenotype, blue, Fig. 6a) and PV (blue, Fig. 6b). These findings suggest synaptic connections of vGluT2+ fibers with neighboring cholinergic and PV neurons in BF, which play a major role in BF function including arousal and sleep/wake regulation, as suggested by previous *in vitro* electrophysiological studies (Yang et al., 2014, 2017; Xu et al., 2015).

**Figure 6.**
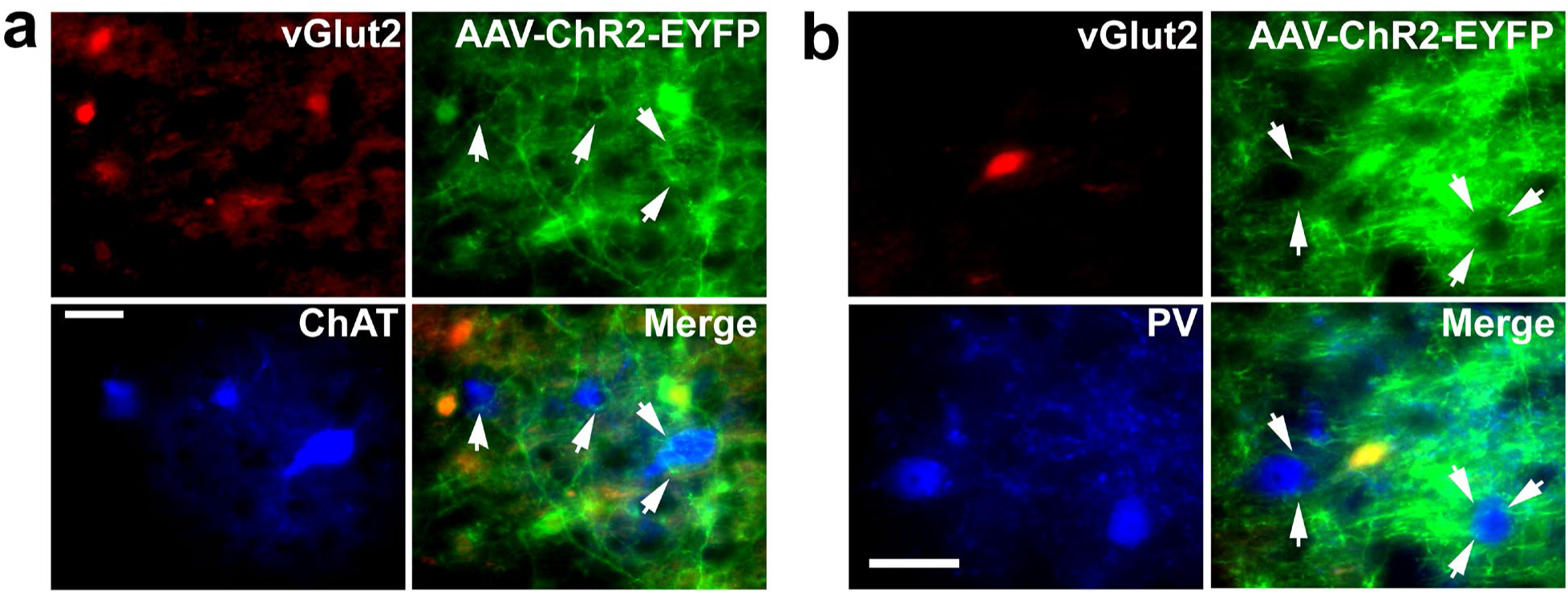
The fibers of BF vGluT2 neurons appose neighboring cholinergic and parvalbumin neurons within BF. a) AAV5-DIO-ChR2-EYFP transduced BF vGluT2+ neuronal fibers (green) appose BF cholinergic neurons (ChAT+, Blue) (indicated by arrowheads) in vGluT2-cre-tdTomato (red) mice. b) Transduced vGluT2+ fibers also appose BF parvalbumin (PV) neurons (indicated by arrowheads). Scale bars = 50 µm.

Next, we examined the projections of BF vGluT2+ neurons outside the transduced area. In the telencephalon, labeling included minor/moderate projections to the cerebral cortex, particularly the medial prefrontal (taenia tecta, prelimbic, and infralimbic cortices) and anterior cingulate cortices (Fig. 7a), as well as the lateral orbital and endopiriform regions (Fig. 7b). Labeled fibers were strongly evident in rostral regions of the BF including the medial septum and horizontal limb of the diagonal band (Fig. 7c), in the olfactory tubercle, as well as in the lateral septum, particularly in its intermediate aspect. Moderate labeling was observed in the amygdala included the basolateral, basomedial, and central nuclei. Transduced fibers were also seen in extended amygdala regions including the bed nucleus of stria terminalis.

**Figure 7.**
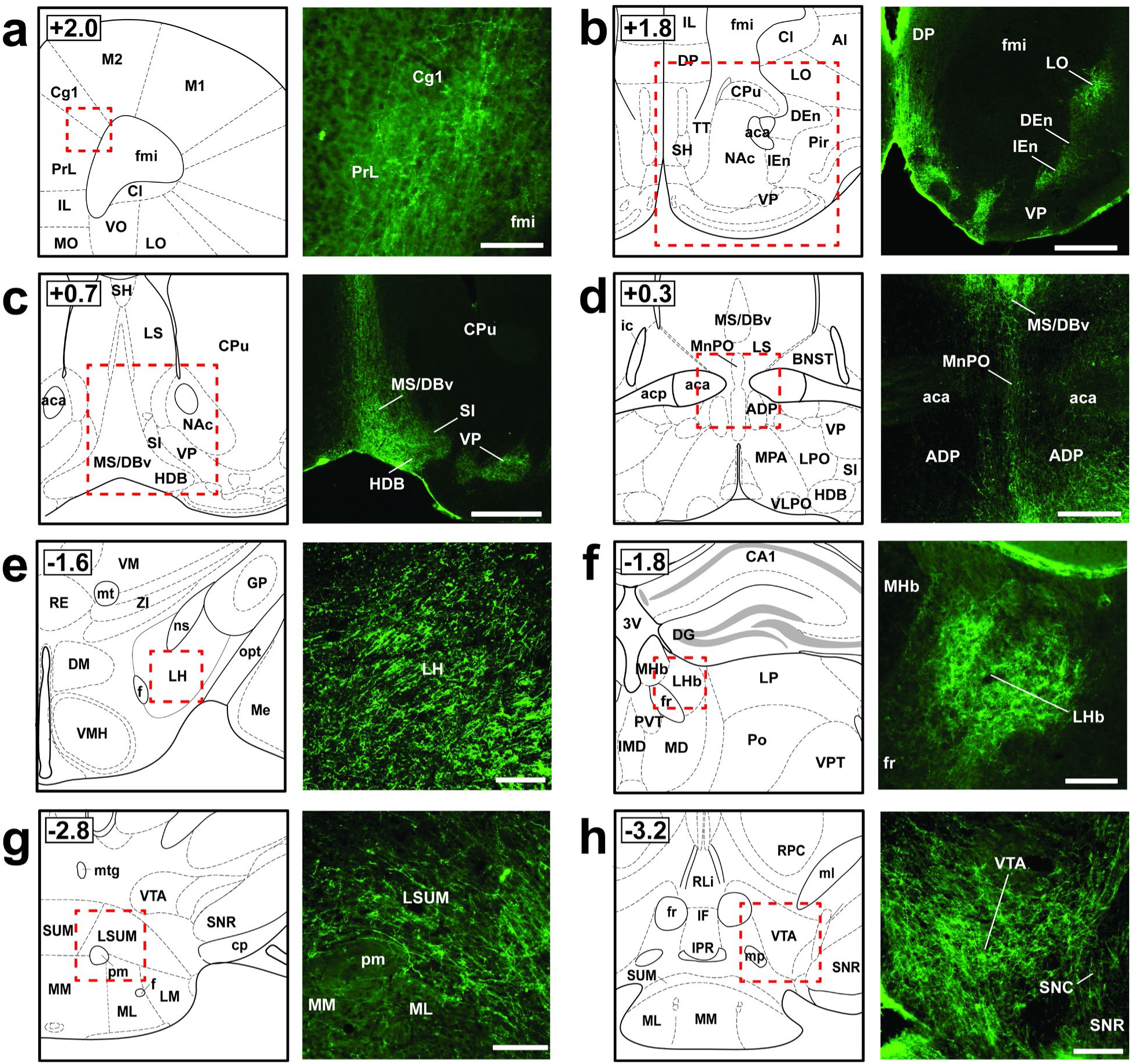
BF vGluT2 fibers were evident in numerous brain regions following AAV5-DIO-ChR2-EYFP injection into BF. Low magnification fluorescent photomicrographs of coronal sections throughout the brain show patterns of labeling produced by AAV5-DIO-ChR2-EYFP injection into BF of vGluT2-cre-tdTomato mice. Moderate/strong EYFP labeled fibers (green) innervate cortical regions including (a) the prelimbic and anterior cingulate cortices; and (b) lateral orbital and endopiriform cortices. Pronounced labeling was seen in (c) the medial septum/diagonal band and lateral septum, as well as areas just caudal including (d) the median preoptic nucleus. Labeling in the hypothalamus included the posterior and lateral hypothalamic regions (e); and strong labeling was noted in the (f) epithalamic lateral habenula. Furthermore, strong fiber labeling was noted in (g) the lateral supramammillary nucleus and neighboring (h) ventral tegmental area of Tsai. See list for abbreviations.

In the diencephalon, labeling was seen throughout much of the hypothalamus, including the arcuate nucleus, dorsomedial hypothalamic nucleus, posterior and lateral hypothalamic regions (Fig. 7e) and lateral supramammillary nuclei (Fig. 7g). Preoptic areas showing intermediate labeling included the median preoptic nucleus (Fig. 7d) and lateral preoptic area. We note the possibility that preoptic labeling may reflect transduction of neighboring non-BF vGluT2+ neurons, particularly in the 2 cases in which viral transduction slightly impinged on hypothalamic/preoptic regions (Fig. 5c). Labeling was largely consistent across all six cases, though, including the four cases in which AAV injections were restricted to BF.

Many thalamic nuclei were labeled following transduction of BF vGluT2 neurons, especially midline and intralaminar nuclei. Intralaminar nuclei which exhibited labeling included the anterior centromedial and posterior parafascicular nuclei. Moderate labeling was also noted in the midline nucleus reuniens, as well as the paratenial, paraventricular, centromedial, mediodorsal, intermediodorsal, anterodorsal, and ventromedial thalamic nuclei. An absence of labeling in the reticular thalamic nucleus was a notable contrast with the prominent fiber labeling seen in this structure after transduction of BF PV neurons with the same viral vector (Kim et al., 2015; Thankachan et al., 2019). Within the epithalamus, a strong labeling of the lateral habenula (Fig. 7f), but not the neighboring medial habenula, was one of the most consistent and striking findings. Projections to the subthalamic zona incerta nucleus were also observed.

BF vGluT2 descending projections to caudal midbrain and brainstem efferent recipients included the substantia nigra pars compacta, ventral tegmental area of Tsai (Fig. 7h), the periaqueductal gray (largely lateral subregions), parabrachial area, nucleus incertus, red nucleus, median and dorsal raphe nuclei, laterodorsal tegmentum, and reticular formation subregions, particularly its isthmic and mesencephalic aspects.

When organized according to functional roles (Fig. 8), our BF vGluT2 circuit mapping revealed numerous efferent projections to brain nuclei that play a crucial role in arousal and vigilance state regulation (Brown et al. 2012), including moderate ascending projections to frontal cortex (cingulate, prefrontal, and orbital cortical regions), moderate/strong projections to forebrain regions including the medial and lateral septum, the lateral and preoptic hypothalamic regions, as well as the supramammillary nuclei (largely its lateral aspect). Minor/moderate projections were also noted to some thalamo-cortical nuclei that are responsible for the cortical activation evident during arousal. Descending projections to arousal-related nuclei included the parabrachial nucleus, select raphe nuclei, tegmental, and reticular formation subregions. Notably, moderate/strong projections were observed in areas involved in hippocampal theta rhythm generation including the ventral tegmental area, dorsal and medial raphe nuclei of the brainstem, suprammamillary nucleus of the hypothalamus, medial septum and horizontal limb of the diagonal band, anteromedial, anterodorsal and reuniens nuclei of the thalamus, hippocampal formation, and select prefrontal cortex regions.

**Figure 8.**
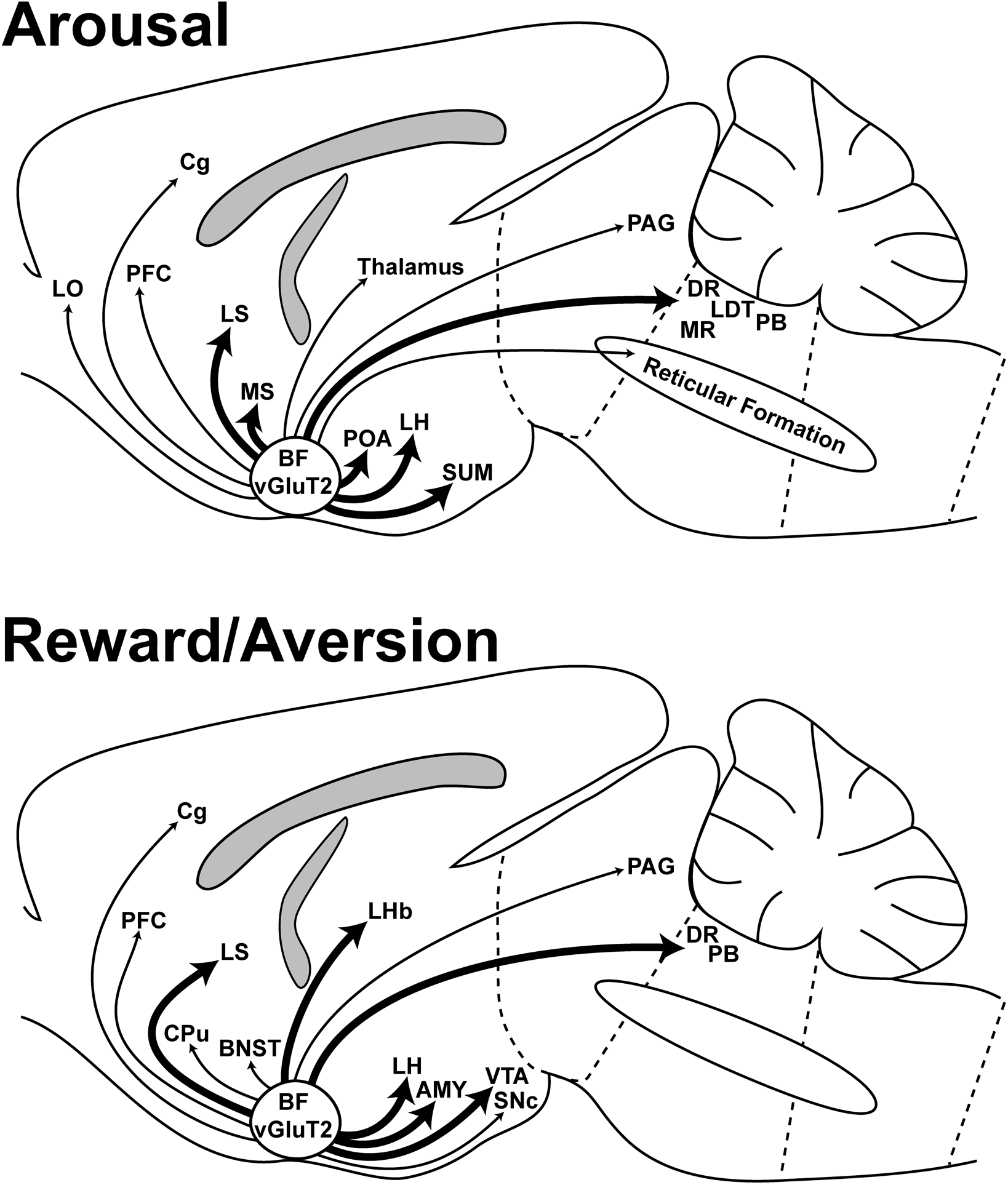
Schematic depiction of BF vGluT2 projections to brain regions involved in arousal and in the response to aversive and rewarding stimuli. *Top:* Strong projections of BF vGluT2 neurons were observed to the following areas involved in arousal regulation and theta rhythm generation: medial and lateral septum, preoptic area, lateral hypothalamus, supramammillary nucleus and dorsal raphe. Less dense projections were observed in frontal cortices, midline thalamus and brainstem arousal nuclei. *Bottom:* Strong projections of BF vGluT2 neurons were observed in the following areas involved in the response to aversive and rewarding stimuli: Lateral septum, lateral habenula, lateral hypothalamus, ventral tegmental area, amygdala and dorsal raphe. Less dense projections were observed in prefrontal/cingulate cortices, BNST, and PAG.

Beyond arousal and theta rhythm regulatory neural circuits, we noted dense fiber concentrations in nuclei involved in the response to aversive and rewarding stimuli (Fig. 8), most notably the lateral habenula, an area which has been closely linked to negative reward predictions and aversive behavior (Tian & Uchida 2015; Lazaridis et al., 2019), as well as the ventral tegmental area and lateral hypothalamus. Minor/moderate projections were also observed in cortical regions implicated in reward neural systems regulation, including frontal cortical regions (infralimbic, prelimbic, anterior cingulate and orbitofrontal cortices). A number of other reward-related nuclei also exhibited moderate/strong labeling, such as the bed nucleus of stria terminalis, interfascicular nucleus, thalamic lateral habenula, caudate putamen, and substantia nigra pars compacta. Moderate labeling was also seen in other regions rich in dopaminergic neurons including zona incerta and A11-14.

### Optogenetic stimulation of BF vGluT2 elicits a marked place aversion

The marked projections of vGluT2 neurons to the lateral habenula and other brain areas involved in reward and aversive behavior led us to investigate the effect of optogenetically activating BF vGluT2 neurons in a place-preference paradigm (Fig. 9). Control experiments targeted two other BF cell types which our immunohistochemical staining experiments indicated are separate from vGluT2-tdTomato neurons; i.e. ChAT and PV neurons. Mice received optogenetic stimulation at a frequency close to their maximal discharge frequency during wakefulness whenever they entered one, randomly chosen side of the chamber. When such stimulation was given bilaterally to vGluT2-cre mice, the mice immediately left that side of the chamber (Video 1, Fig. 9), a dramatic effect which was repeated many times over the course of the 30 min session in all 4 mice tested. Overall, mice spent only 152 ± 49 s on the stimulated side versus 1102 ± 157 s on the unstimulated side, resulting in a place-preference ratio of 0.16 ± 0.06 (t=4.60, P=0.0193, paired t-test), indicating a strong place aversion. Surprisingly, when mice were returned to the chamber on the following day, in the absence of optogenetic stimulation this place aversion was not maintained (Fig. 9, place-preference ratio of 0.83 ± 0.13, n.s.).

**Figure 9.**
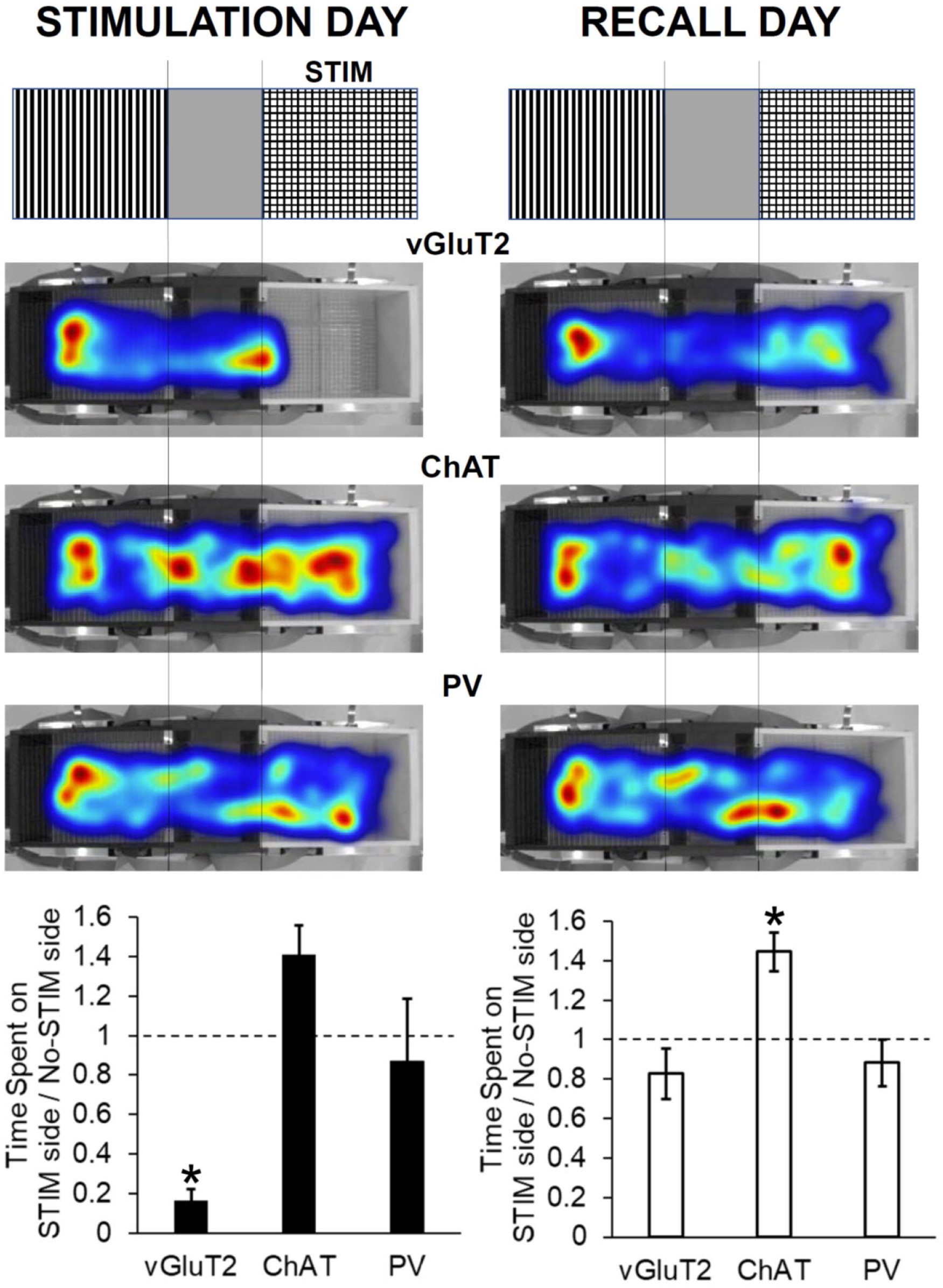
Optogenetic stimulation of BF vGluT2 neurons elicits a marked place aversion. *Left:* BF vGluT2, cholinergic (ChAT) or parvalbumin (PV) neurons were briefly stimulated at a frequency close to their maximal discharge rate during wakefulness whenever they entered one, randomly chosen side of a three-part chamber, depicted schematically in the top part of the figure. Heatmaps showing the location of the midpoint of the mice (middle part of the figure) clearly show that the mice with stimulation of vGluT2 mice neurons avoid the area where optogenetic stimulation is given, whereas stimulation of ChAT or PV neurons does not. Quantification of these results (bottom) showed that stimulation of vGluT2 neurons results in a significant (p<0.05) place aversion (n = 4). Stimulation of cholinergic neurons (n = 3) tends to produce a place preference whereas stimulation of PV neurons does not lead to a place preference or a place aversion (n = 3). *Right:* Returning the mice to the chamber one day later in the absence of optogenetic stimulation reveals that BF vGluT2 mice do not retain a place avoidance whereas BF ChAT mice retain a significant place preference for the side where optical stimulation was given. PV mice do not show a preference.

Very different results were obtained when we performed the same type of experiment for BF cholinergic and parvalbumin neurons. Bilateral optogenetic stimulation of cholinergic neurons in ChAT-cre mice led to a trend-level place preference instead of a place aversion (Video 2, Fig. 9, Place preference ratio 1.41 ± 0.15, n = 3, t=3.37, p=0.078) which was maintained on the next day, showing a significant place preference in the absence of optogenetic stimulation (1.45 ± 0.10, t=6.39, p=0.024). Bilateral optogenetic stimulation of BF PV neurons in PV-cre mice did not cause a place preference or a place aversion (Video 3, Fig. 9) either on the stimulation day (place preference ratio 0.87 ± 0.31, n=3, n.s.) or on the next day (0.88 ± 0.12, n.s.). In addition to providing a control for our experiments with vGluT2 neurons, these experiments suggest that the effect of BF vGluT2 stimulation in producing a place aversion is not due to local excitatory effects of BF vGluT2 neuronal stimulation on neighboring cholinergic and PV neurons.

## DISCUSSION

The main findings reported in this study are as follows: (i) vGluT2-tdTomato is not expressed in BF cholinergic and GABAergic neurons; (ii) BF vGluT2-tdTomato neurons are almost all small or medium sized neurons and are less dense in VP of BF; (iii) Major subsets of BF vGluT2-tdTomato neurons co-express Calb and Calr, but rarely PV; (iv) BF vGluT2+ neurons project within BF to cholinergic and PV neurons and outside BF to multiple areas involved in regulation of arousal and the response to aversive and rewarding stimuli; and (v) Optogenetic stimulation experiments showed that stimulation of these neurons elicits strong avoidance behavior. In the subsequent sections we discuss each of these findings and their importance.

### vGluT2-cre-tdTomato mice can be used to reliably identify BF glutamatergic neurons

Identification of glutamatergic neurons has proven challenging due to the lack of specific methods to identify them. Although staining for phosphate-activated glutaminase (Manns et al., 2001; Gritti et al., 2006) or glutamate itself (Ottersen and Storm-Mathisen, 1985) gave some hints about the distribution and function of BF glutamatergic neurons, these immunohistochemical stains are not specific for glutamatergic neurons (Kaneko and Mizuno, 1988; Akiyama et al., 1990; Laake et al., 1999), likely due to the involvement of glutamate in cellular metabolism and as a precursor for GABA synthesis. The identification of the genes for the vesicular glutamate transporters allowed glutamatergic neurons to be unequivocally identified using *in situ* hybridization for the first time (Fremeau et al., 2001, 2004; Herzog et al., 2001; Hur and Zaborszky, 2005). However, *in situ* hybridization is not easy to combine with *in vitro* electrophysiological recordings, and immunohistochemical staining for vGluT2 does not reliably label cell bodies (Henny and Jones, 2006, 2008), hampering the identification of the cellular properties of glutamatergic neurons. In some studies, single-PCR for vGluT mRNA has been used to identify glutamatergic neurons (Sotty et al., 2003). However, this technique has a tendency for false-positives and does not allow targeting of putative glutamatergic neurons prior to *in vitro* recording. Therefore, the development of mice with Cre recombinase expressed under the control of the vGluT2 promoter was a considerable advance (Vong et al., 2011).

In order to use vGluT2-tdTomato mice to characterize BF glutamatergic neurons *in vitro* and/or for optogenetic or pharmacogenetic experiments *in vivo*, it is important to verify that ectopic expression does not occur in cholinergic and GABAergic neurons, the other two major subtypes of BF neurons. Our results suggested that there was no or very little ectopic expression in the BF, consistent with results in vGluT2-cre mice in other subcortical regions (Vong et al., 2011) and with experiments using *in situ* hybridization to confirm selective expression (Anaclet et al, 2015; Xu et al., 2015). In particular, we observed no colocalization with ChAT, a selective marker for cholinergic neurons. Similarly, no colocalization with GFP was observed in vGluT2-tdTomato/GAD67-GFP knock-in mice, with GFP selectively expressed in GABAergic neurons (McKenna et al., 2013). Thus, vGluT2-cre-Tomato mice can be used to study the major subpopulation of BF glutamatergic neurons.

### BF vGluT2-tdTomato neurons in the mouse do not label for ChAT, indicating lack of co-expression with cholinergic neurons

Co-localization of glutamate with acetylcholine in BF had been previously suggested (Forloni et al., 1987), including co-release at fiber terminals (Docherty et al., 1987; Allen et al., 2006), although a follow-up investigation suggested a much smaller percentage of co-localization of glutamate and acetylcholine in BF neuronal soma (Szerb and Fine, 1990). Immunohistochemical staining for vGluT2 and the vesicular acetylcholine transporter (VAChT) combined with anterograde tracing of efferent projections emanating from BF in the rat did not reveal any co-localization in varicosities in the cortex or in the lateral hypothalamus, brain regions in which both neurotransmitter markers were expressed (Henny and Jones, 2006; 2008). As reported here, vGluT2-tdTomato neurons had a different distribution and size profile than cholinergic neurons. Our previous *in vitro* study also suggest that they have distinct intrinsic membrane properties (Yang et al., 2017). Thus, several convergent lines of evidence suggest that cholinergic neurons do not express vGluT2.

A subpopulation of BF cholinergic neurons which largely lack the p75 neurotrophin receptor expresses vGluT3, with highest levels of co-localization in the ventral pallidum (Nickerson Poulin et al., 2006). Using retrograde tracer techniques, this unique subset of BF neurons was shown to project to the basolateral amygdala. ChAT and vGluT3 co-localization was analyzed in a number of areas known to be strongly innervated by the cholinergic component of BF, and co-localization in fibers was noted only in the amygdala, but not other brain regions. Also, vGluT3 was not detected in axonal terminals of BF cholinergic projections to other brain regions outside of the amygdala, including the neocortex and lateral hypothalamus (Henny and Jones, 2006, 2008). Furthermore, electrophysiological studies of the BF cholinergic projection to the amygdala did not reveal a glutamatergic component (Unal et al., 2015). Overall, these findings further suggest that BF cholinergic neurons do not release glutamate as a co-transmitter at their distal targets, although a local autoregulatory role involving vGluT3 cannot be ruled out.

### BF vGluT2-tdTomato neurons in the mouse do not co-localize with GAD67-GFP

Once thought infeasible, co-release of glutamate and GABA has now been demonstrated at several synapses in the mammalian nervous system, in particular at entopeduncular and ventral tegmental projections to the habenula (Root et al., 2014; Shabel et al., 2014) and in the anteroventral periventricular nucleus of the hypothalamus (Ottem et al., 2004). Switching of the relative contribution of glutamatergic and GABAergic neurotransmission at these synapses is a newly identified mechanism of neural plasticity which allows a change in the sign of the synapse (excitatory or inhibitory) in response to changing behavioral needs (Shabel et al., 2014). Given these findings, we carefully examined whether there was co-expression of tdTomato and GFP in vGluT2-tdTomato/GAD67-GFP crossed mice. Our results demonstrate that vGluT2 is not co-expressed in GABAergic (GAD67+) neurons in BF. Taken together, our findings suggest that ChAT+, GAD67+ and vGluT2-tdTomato neurons represent three largely non-overlapping neuronal BF subpopulations, as also suggested by findings from other groups (Henny and Jones, 2008; Hassani et al., 2009; Anaclet et al., 2015; Xu et al., 2015). We note, however, that these findings do not exclude the possibility that some cholinergic neurons may co-express GAD65 and release GABA at their cortical targets (Saunders et al., 2015).

### vGluT2-tdTomato neurons represent a subpopulation of small to mid-sized neurons scattered throughout BF

vGluT2-tdTomato neurons were located throughout BF, similar to previous *in situ* mapping studies of the rat BF (Fremeau et al., 2001; Herzog et al., 2001; Lin et al., 2003; Hur and Zaborszky, 2005), as well as investigations employing the vGluT2-IRES-cre mouse model used here (Anaclet et al., 2015; Xu et al., 2015; Do et al., 2016; Tooley et al., 2018). However, a detailed mapping and analysis of vGluT2 neurons in mice had not previously been reported. The density of vGluT2-tdTomato neurons in the VP subnuclei was significantly decreased compared to other BF subregions, similar to previous studies in the rat, where the vGluT2+ neuronal population was noted as “scarce” in VP (Hur and Zaborszky, 2005). Glutamatergic neurons in the rat BF were qualitatively described as small-to-medium sized, similar to the measures of vGluT2-tdTomato long-axis diameters reported here, as well as in our *in vitro* investigations employing the same vGluT2-tdTomato mouse model (Yang et al., 2017). Overall, vGluT2-tdTomato neurons are smaller in size than mouse BF GABAergic and PV+ neurons (McKenna et al., 2013). Our *in vitro* and anatomical investigations (McKenna et al., 2013; Yang et al., 2014, 2017) therefore suggest that, in the mouse, the average BF long-axis diameter is largest in cholinergic neurons, followed by GABAergic neurons, and the smallest BF neurons are of the glutamatergic vGluT2+ neuronal phenotype.

### Major subsets of BF vGluT2-tdTomato neurons in the mouse express the calcium binding proteins Calb and Calr, but not PV

The calcium binding proteins Calb, Calr, and PV are commonly used as markers for functionally distinct subsets of GABAergic interneurons in the cortex, hippocampus and striatum (Celio, 1986; Kawaguchi and Kubota, 1993; Kubota et al., 1993). However, this association of calcium binding proteins with GABAergic interneurons does not always hold in subcortical regions. Here we found that major subsets of BF vGluT2-tdTomato neurons express Calb and Calr, but few express PV, which appears to a be a selective marker for a subset of very fast-firing BF GABAergic projection neurons (McKenna et al., 2013) involved in the control of cortical gamma band oscillations (Kim et al., 2015) and arousals from sleep (McKenna et al., 2020).

*Calb.* Calb+ neurons have been previously described in the rat BF (Riedel et al., 2002; Gritti et al., 2003), including co-localization with GABAergic and cholinergic populations. One study in the rat proposed that ∼40% of BF Calb+ neurons co-localized with phosphate-activated glutaminase (Gritti et al., 2003) but this enzyme is not a selective marker for glutamatergic neurons. Thus, our study is the first to definitively identify a subpopulation of BF glutamatergic (vGluT2+) neurons that co-express Calb.

*Calr.* In addition to PV and Calb, Calr has also been previously reported as a calcium binding protein localized in a subset of neurons distributed throughout BF in the rat (Kiss et al., 1997; Riedel et al., 2002; Gritti et al., 2003). Calr+ neurons in the rostral BF, including the medial septum and horizontal limb (both HDB and VDB), were not cholinergic and did not co-localize with PV or Calb (Kiss et al., 1997). Calr was reported to be present in a subset of non-cholinergic, non-GABAergic neurons which do not project to the cerebral cortex (Gritti et al., 2003). A significant proportion of Calr+ fibers in the entorhinal cortex were identified as glutamatergic (Wouterlood et al., 2000), although it was not determined if BF was a source of this projection. Overall, growing evidence supports the suggestion that Calr is present in a subpopulation of BF glutamatergic neurons, as we directly demonstrate here.

### Intra-BF glutamatergic projections to neighboring PV+ and cholinergic BF subpopulation

Examination of vGluT2+ fibers/varicosities in BF indicated that vGluT2+ neurons appose local cholinergic and PV+ neurons. Early investigations employing *in situ* hybridization suggested that vGluT2+ neurons in the neighboring MS/DBV provide input to local PV+ neurons that project to the hippocampus and play a major role in promotion of hippocampal theta rhythmicity (Wu et al., 2003; Hajszan et al., 2004). By combining optogenetics and *in vitro* patch clamp recordings, a recent study described involvement of local MS/DBV glutamatergic projections to neighboring GABAergic and, to a lesser extent, cholinergic neurons, as part of the intra-septal mechanism responsible for hippocampal theta generation (Robinson et al., 2016). Using the vGluT2-IRES-cre mouse model, *in vitro* electrophysiology and optogenetic experiments demonstrated local BF glutamatergic excitatory inputs to PV+ and cholinergic neurons (Xu et al., 2015). While definitive evidence of synaptic contacts requires electron microscopic analysis, collectively these light microscopy and *in vitro* electrophysiology and optogenetic experiments provide evidence for excitatory inputs from BF vGluT2 neurons to local PV and cholinergic neurons. *In vivo* optogenetic experiments suggested that strong synchronous activation of BF vGluT2 neurons promotes arousal (Xu et al., 2015), which may involve both the intra-BF connections as well as the extra-BF projections to other arousal promoting regions described here and in previous studies (Anaclet et al., 2015; Do et al., 2016; Agostinelli et al., 2017, 2019).

### Extra-BF glutamatergic projections to arousal-related brain regions

Our BF vGluT2 specific anterograde mapping revealed fiber labeling in brain regions previously described to play a crucial role in sleep-wake control and cortical activation (Fig. 8). The BF is a key forebrain node in the brain circuitry which promotes wakefulness and promotes high-frequency cortical activity typical of conscious states (Brown et al., 2012). However, the role of individual BF cell-types is still being uncovered. A previous study showed that strong optogenetic stimulation of BF vGluT2 neurons was highly potent in inducing prolonged periods of wakefulness (Xu et al., 2015). Weaker chemogenetic activation of the BF vGluT2 neuronal population was less potent in promoting arousal than activation of GABAergic neurons (Anaclet et al., 2015), possibly due to the relatively hyperpolarized resting membrane potential of BF vGluT2 neurons (Yang et al., 2017) compared to GABAergic neurons (McKenna et al., 2013). In addition to the local effects on other BF subtypes described above, our anterograde tracing experiments here suggest pathways by which strong activation of BF vGluT2 neurons may promote arousal.

Projections to the cortex were relatively sparse/moderate, including labeling in prefrontal, orbital, and endopiriform cortices, consistent with previous tracing studies (Hur and Zaborszky, 2005; Do et al., 2016). In particular, our findings confirm previous retrograde tracer studies that revealed a sparse projection from BF glutamatergic neurons to the medial prefrontal cortex (mPFC) in the rat, in which the glutamatergic component of the total BF-mPFC projection was estimated to be ∼5% (Hur and Zaborszky, 2005). Manns et al. (2001) reported a higher estimate of glutamatergic contribution to the BF efferent projection to the entorhinal cortex, employing co-labeling of BF retrograde labeled neurons with immunohistochemical detection of phosphate-activated glutaminase. Again, immunolabeling for this enzyme may also detect GABAergic neuronal processes, contributing to a possible overestimation of BF glutamatergic projections. Considering this evidence together, BF vGluT2 output is a relatively small component of the overall direct BF-cortical projection.

Considering the relatively small direct BF glutamatergic projection to cortex, projections of BF glutamatergic neurons to other subcortical regions may be responsible for the wakefulness-promoting effects associated with optogenetic stimulation of these neurons (Xu et al., 2015). In the forebrain, moderate/strong fiber labeling was seen in the bed nucleus of stria terminalis, which has been recently described as playing a role in cortical activation (Kodani et al., 2017), as well as in the supramammillary region which promotes wakefulness and theta activity (Vertes 2015; Pedersen et al., 2017). Dense fiber labeling was also observed in a key theta-rhythm generating area, the MS/DBV (Vertes and Kocsis, 1997).

Fiber labeling was moderate to strong in the lateral hypothalamic area, where orexin/hypocretin neurons that promote consolidated wakefulness are located, confirming projections reported by other investigators (Henny and Jones, 2006; Agostinelli and Scammell 2017, 2019; Do et al., 2017). Previous work in rats (Henny & Jones 2006) and mice (Agostinelli et al., 2017) showed that BF vGluT2 fiber terminals appose orexin/hypocretin neurons and release glutamate (Agostinelli et al., 2017). Thus, this projection is one likely candidate to explain the increased wakefulness produced by stimulation of BF vGluT2 neurons. Moderate labeling was also observed in numerous thalamic regions, largely localized to select midline and neighboring thalamo-cortical nuclei previously described as essential for cortical activation (anterodorsal, mediodorsal, centromedial, intermediodorsal, and the reuniens/rhomboid nuclei) (Steriade et al., 1993; Van der Werf et al., 2002; Gent et al., 2018).

Although largely implicated in reward system processing recent evidence suggests that the ventral tegmental area of Tsai (VTA) may also play a role in arousal and cortical activation (Eban-Rothschild et al., 2016; Oishi et al., 2017; Sun et al., 2017; Yu et al., 2019) and defensive responses (Barbano et al., 2020). We found particularly strong fiber labeling here in VTA, one of the strongest BF vGluT2+ projections. Sparse/moderate labeling was observed in a number of classic posterior midbrain and brainstem arousal nuclei, including the laterodorsal and pedunculopontine tegmental regions, dorsal and median raphe, and the reticular formation, particularly noted in its isthmic and mesencephalic subregions. Moderate fiber labeling was also evident in the parabrachial nucleus, thought to be a key brainstem nucleus responsible for cortical activation (Fuller et al., 2011).

### BF vGluT2+ neurons project to multiple regions involved in processing aversive and rewarding stimuli

As depicted in Figure 8b, our anterograde mapping studies also revealed BF vGluT2 input to many nuclei involved in processing aversive and rewarding stimuli, including VTA. Glutamatergic inputs to both dopaminergic and non-dopaminergic VTA neurons play a key role in regulation of VTA firing (Lammel et al., 2014), suggested by studies employing glutamatergic agonist infusion (Grace and Bunney, 1984; Johnson et al., 1992; Chergui et al., 1993) and regulate self-stimulation and the effects of drugs of abuse, including cocaine and opiates (Panagis and Kastellakis, 2002; Harris and Aston Jones, 2003; Harris et al., 2004). A previous investigation in the rat determined that a small portion of the BF projection to VTA from the VP region of BF is vGluT2+ (Geisler et al., 2007). However, as we show here, VP has the lowest density of BF vGluT2-tdTomato neurons. Thus, other BF subregions may have more prominent roles. Input from VP to dopaminergic, glutamatergic, and GABAergic VTA neurons has been demonstrated (Faget et al., 2016), but the relative importance of the projection to each neuronal class remains to be determined. A recent study concluded that VP glutamatergic input to VTA GABAergic VTA neurons elicits place aversion (Tooley et al., 2018), consistent with our *in vivo* optogenetic findings of a strong place aversion elicited by stimulation of a larger proportion of BF vGluT2+ neurons. However, it is likely that other projections of BF vGluT2+ neurons also are important in eliciting place aversion.

In addition to the prominent BF vGluT2+ projection to VTA, some of the strongest fiber labeling was evident in the lateral habenula, which plays a crucial role in negative reward predictions (Tian and Uchida, 2015; Baker et al., 2016) and depressive behavior (Proulx et al., 2014; Browne et al., 2018). Recent studies have suggested that VP glutamatergic projections to the lateral habenula and VTA may promote behavioral response to stress or punishment avoidance (Tooley et al., 2018; Wulff et al., 2019), consistent with the response we observed here *in vivo*.

Besides the VTA and lateral habenula, moderate to strong inputs were observed to other nuclei important in processing aversive and rewarding stimuli (Figure 7b) such as the lateral septum (Zernig and Pinheiro, 2015; Tsanov, 2018), amygdala (Baxter and Murray, 2002; Murray, 2007; Janak and Tye, 2015), lateral hypothalamus (Aston-Jones et al., 2010; Castro et al., 2015; Tyree and de Lecea, 2017) and dorsal raphe (Ikemoto and Bonci, 2014; Luo et al., 2015; Li et al., 2016). Cortical projections to the medial prefrontal and orbitiofrontal regions are also consistent with a role in processing of aversive/rewarding stimuli (Weinberger et al, 1988; Lammel et al., 2014; Boot et al., 2017).

### Optogenetic activation of BF vGluT2 neurons elicits a striking place avoidance response

The prominent projections of BF vGluT2 neurons to brain areas involved in processing aversive and rewarding stimuli led us to test the effect of activating them *in vivo* in a place preference paradigm. Bilateral optogenetic activation of these neurons led to a striking and highly consistent avoidance response which was markedly different from the response in control experiments which targeted BF cholinergic or PV neurons. The different responses in cholinergic and PV neurons also suggest that the effect of stimulating BF vGluT2 neurons is not mediated by the local connections within BF. Instead, our anterograde tracing results presented here together with prior literature suggest that the avoidance response elicited by optogenetic stimulation of BF vGluT2 neurons is most likely mediated by a coordinated activation of neurons in the lateral habenula, lateral hypothalamus and VTA (Knowland et al., 2017; Tooley et al., 2018; Laziridis et al., 2019; Patel et al., 2019; Barbano et al., 2020).

The avoidance response we observed is consistent with a growing body of work implicating BF vGluT2 neurons in the response to aversive stimuli and in constraining reward seeking (Faget et al., 2016; Tooley et al., 2018; Patel et al., 2019). Surprisingly, in our experiments this avoidance response was not maintained one day later. Thus, BF vGluT2 neurons are involved in immediate avoidance responses but one 30 min session of optogenetic stimulation was not sufficient to develop a learned avoidance response. The signals which activate BF vGluT2 neurons to cause avoidance behavior are not fully understood but an interesting recent study identified an excitatory population of BF neurons that are activated by food odor-related stimuli and drive selective avoidance of food with unpleasant odors, in part via projections to the lateral hypothalamus (Patel et al., 2019).

Optogenetic stimulation of cholinergic neurons elicited a place preference, opposite to the place avoidance we observed with stimulation of vGluT2 neurons. Interestingly, our *in vitro* recordings (Yang et al., 2017) identified a long-lasting inhibitory response to the cholinergic agonist, carbachol, in a subset of vGluT2 neurons located in the ventromedial BF, the same BF subregion where Patel and colleagues (2019) recorded vGluT2 neurons which responded to aversive food odors. Thus, these two groups of BF neurons may act in an antagonistic manner.

### Conclusion

This study provides a comprehensive description of vGluT2+ neurons in the mouse BF. vGluT-tdTomato neurons are distinct from neighboring cholinergic and GABAergic BF populations. The calcium binding proteins Calb and Calr are co-localized in subsets of BF vGluT2-tdTomato neurons indicating distinct functional subtypes. Our cell-specific anterograde investigation of BF vGluT2+ projections, together with our *in vivo* optogenetic experiments and previous literature proposing a prominent role in promoting wakefulness (Xu et al., 2015), suggest that these neurons are uniquely positioned to promote arousal and avoidance behavior in response to aversive stimuli. Several psychiatric disorders such as post-traumatic stress disorder, anxiety disorders and alcohol use disorder are associated with both increased arousal and enhanced aversive responses (Sharma et al., 2010; Lind et al., 2020). Thus, in the future it will be of interest to investigate a potential role for BF vGluT2 neurons in animal models of these disorders.

## Role of authors

JTM performed, oversaw, designed and analyzed anatomical experiments to validate vGluT2-tdTomato mice, described the localization, size and colocalization with calcium-binding proteins and performed experiments analyzing the anterograde projections. JTM. also supervised other study personnel performing anatomical experiments, generated figures 1-8 and wrote the first draft of the manuscript together with REB. CY performed, designed and analyzed the optogenetic place preference experiments and generated figure 9 and the associated videos. TB mapped anterograde projections of BF vGluT2 neurons together with JTM. MA-C, MG, AH and JM analyzed anatomical data. SW performed immunohistochemical staining, transcardial perfusions and tissue mounting. JMM assisted with the optogenetic place preference experiments and provided comments on the manuscript. RB provided mice for the place preference experiments, participated in the discussion of the results, and provided comments on the manuscript. REB conceived the study, obtained funding and regulatory approvals, designed experiments, analyzed and interpreted data and wrote and edited the manuscript.

## Table of abbreviations

3V: Third ventricle
A11: A11 dopamine cells
A12: A12 dopamine cells
A13: A13 dopamine cells
A14: A14 dopamine cells
AAV: Adeno-associated virus
aca: Anterior commissure, anterior
acp: Anterior commissure, posterior
ADP: Anterodorsal preoptic nucleus
AI: Agranular insular cortex
AMY: Amygdala
BF: Basal forebrain
BNST: Bed nucleus of stria terminalis
CA1: Hippocampal field CA1
Calb: Calbindin Calr Calretinin
CPu: Caudate Putamen
cc: Corpus callosum
Cg: Cingulate cortex
Cg1: Cingulate cortex, area 1
ChAT: Choline acetyl transferase
ChR2: Channelrhodopsin 2
Cl: Claustrum
cp: Cerebral peduncle
CTX: Cotex
DBv: Vertical limb of the diagonal band
DEn: Endopiriform nucleus, dorsal
DG: Dentate gyrus
DM: Dorsomedial hypothalamic nucleus
DP: Dorsal peduncular cortex
DR: Dorsal raphe nucleus
EYFP: Enhanced yellow fluorescent protein
f: Fornix
fmi: forceps minor of the corpus callosum
fr: Fascilculus retrofelxus
GAD67: Glutamic acid decarboxylase-67
GFP: Green fluorescent protein
GP: Globus pallidus
HDB: horizontal limb of the diagonal band
ic: Internal capsule
IEn: Endipiriform cortex, intermediate
IF: Interfascicular nucleus
IL: Infralimbic cortex
IMD: Intermediodorsal thalamic nucleus
LH: Lateral hypothalamic area
IPR: Interpeduncular nulceus, rostral
LDT: Laterodorsal tegmental nucleus
LHb: Lateral habenula
LM: Lateral mammillary nucleus
LO: Lateral orbital cortex
LP: Lateral posterior thalamic nucleus
LPO: Lateral preoptic area
LS: Lateral septum
LSUM: Supramammillary nucleus, lateral
M1: Primary motor cortex
M2: Secondary motor cortex
MCPO: Magnocellular preoptic nucleus
MD: Mediodorsal thalamic nucleus
Me: Medial amygdaloid nucleus
MHb: Medial habenula
ml: medial lemniscus
ML: Medial mammillary nucleus, lateral
MM: Medial mammillary nucleus, medial
MnPO: Median preoptic nucleus
MO: Medial orbital cortex
mp: Mammillary peduncle
MPA: Medial preoptic area
mPFC: Medial prefrontal cortex
MR: Medial raphe nucleus
MS: Medial septum
mt: Mammillothalamic tract
mtg: Mammillotegmental tract
NAc: Nucleus accumbens
ns: Nigrostriatal bundle
opt: Optic tract
PAG: Periaqueductal gray
PB: Parabrachial nucleus
PFC: Prefrontal cortex
Pir: Piriform cortex
Pm: Principal mammillary tract
PN: Paranigral nucleus
Po: Posterior thalamic nuclear group
PrL: Prelimbic cortex
PV: Parvalbumin
PVT: Paraventricular thalamic nucleus
RE: Nucleus reuniens
RLi: Rotral linear raphe nucleus
RPC: Red nucleus, parvicellular
SEM: Standard error of the mean
SH: Septohippocampal nucleus
SI: Substantia innomminata
SNC: Substantia nigra, compacta
SNR: Substantia nigra, reticulate
SUM: Supramammillary nucleus
TT: Tenia tecta
Tu: Olfactory tubercle
vGluT2: Vesicular glutamate transporter 2
VLPO: Ventrolateral preoptic nucleus
VM: Ventromedial thalamic nucleus
VMH: Ventromedial hypothalamic nucleus
VO: Ventral orbital cortex
VP: Ventral pallidum
VPT: Ventral posterior thalamic nuclei
VTA: Ventral tegmental area of Tsai
VTM: Ventral tubermammillary nucleus
ZI: Zona incerta

## Notes

**Funding:** This work was supported by United States Veterans Administration Biomedical Laboratory Research and Development Service Merit Awards I01 BX004673, BX001356, I01 BX001404, I01 BX002774, I01 BX004500 and United States National Institute of Health support from NINDS R21 NS093000, NIMH R01 MH039683, NHLBI HL095491, R03-MH107650 and by SURE fellowships from Stonehill College. JTM, JMM, RB and REB are Research Health Scientists at VA Boston Healthcare System, West Roxbury, MA. The contents of this work do not represent the views of the U.S. Department of Veterans Affairs or the United States Government.

**Conflicts of interest:** No conflicts of interest have been identified for any of the authors. JTM received partial salary compensation and funding from Merck MISP (Merck Investigator Sponsored Programs) but has no conflict of interest with this work.

### Competing Interest Statement

The authors have declared no competing interest.

